# Ribosomal frameshifting is a carefully tuned modifier of proteins

**DOI:** 10.1101/2025.10.02.679949

**Authors:** Jonas Poehls, Cedric Landerer, Doris Richter, Maria Luisa Romero Romero, Anna Shevchenko, Andrej Shevchenko, Agnes Toth-Petroczy

## Abstract

Ribosomal frameshifting, occurring during protein translation, is one of the most consequential recoding events, as it leads to large changes to the protein sequence. However, beyond a few well-characterized cases, its prevalence and consequences across organisms remain poorly understood. To address this, we modeled and experimentally tested frameshifting and systematically characterized its evolutionary impact. We developed SLIPPERRS, a mechanistic model that identifies “slippery” frameshift sites achieving ∼70% accuracy. Building on SLIPPERS’ predictions and on our mass spectrometry and fluorescence-based assay, we characterized 165 short mRNA sequences. We identified dozens of novel sites with frameshift probabilities that often exceed those of known programmed frameshift cases, and found evidence for both translocation and decoding mechanisms. In *Gammaproteobacteria*, an average of ∼100 genes per species harbor frameshifting sites, which alter proteins by truncating, extending, and fusing canonical proteins. We detect selection for shorter and less disordered frameshifted proteins to mitigate their deleterious impact. We found negative correlation between GC-content and the occurrence of frameshifting sites. In GC-rich bacteria, where frameshift products are expected to be particularly long and disordered, there is selection for cryptic stop codons. While frameshifting is prevalent and expands protein diversity, it is carefully tuned to avoid its harmful consequences.

## INTRODUCTION

Translation of mRNA into proteins, a process essential to life, is prone to a variety of errors, collectively referred to as phenotypic mutations^1^. One of the most significant of these errors is ribosomal frameshifting (FS), where the reading ‘shifts’ from one frame to another. Ribosomal frameshifting, like all phenotypic mutations, ‘recodes’ the mRNA and leads to a new protein sequence. This recoding is stochastic: At a given mRNA position under given conditions, a frameshift will happen with a certain probability. The probability of frameshifting depends on several factors: The main element stimulating FS is a ‘slippery site’, a short mRNA sequence (usually 4-10 nucleotides^2–6)^ allowing the binding of a tRNA in several frames^3,6–9^. In addition to the rebinding ability of tRNAs already positioned within the ribosome, frameshifting is thermodynamically controlled by competition among the cellular tRNAs to bind to the A-site^3,10–12^. Frameshifting may happen during either of two phases of translation elongation: i) during decoding, when a tRNA arrives at the ribosome, it may directly bind in a shifted frame^6,11,12^; ii) during the later translocation stage, when two tRNAs move within the ribosome from P- and A-site to E-site and P-site, they can ‘slip’ and bind to a new frame^7,11^. If a slippery site is combined with additional stimulatory elements, such as a stable mRNA secondary structure 3’ of the slippery site, or a 5’ Shine-Dalgarno-like sequence, the frameshift probability can become very high, reaching 50% or more in some cases.

Frameshifts with high probability and an adaptive function are called ‘programmed ribosomal frameshifting’ (PRF). Numerous instances of PRF are known among viruses^5,13,14^, eukaryotes^15–21^, and prokaryotes^22–25^. Examples of functional PRFs include the regulation of protein expression via a frameshift-modulating molecule, as is the case for the bacterial release factor 2^24^ or the eukaryotic antizyme^16^, the control of relative expression levels of two proteins, commonly used by viruses^5,14,26^, and modularity, where two related proteins are produced from the same mRNA (e.g. *copA* and *dnaX* in *E. coli*^23,27^ and *IDP* in yeast^18^). Thus, programmed ribosomal frameshifting serves as a mechanism to regulate and expand proteome diversity.

Apart from PRF, frameshifting is considered rare, occurring at rates of 10^-5^ to 10^-3^, based on studies of leaky frameshift mutants^28,29^ and disrupted PRF signals^13,30^. However, recent studies show that frameshifting probabilities can reach above 1% at inconspicuous sites without evidence of function or a conserved FS signal, both in eukaryotes^31^ and prokaryotes^4^. This suggests effective frameshift sites may be widespread in genomes, with the potential to alter the repertoire of proteins produced by the cell. These new proteins may gain function that is beneficial or deleterious. The production of non-canonical proteins which remain non-functional or misfold/aggregate will waste resources and may be toxic. The rate and type of translation errors are likely under stringent selection at the protein and residue level, as they influence organismal fitness^32,33^.

To fully understand the impact of frameshifting on the proteome and its evolutionary impact, it is essential to identify the location of the frameshift sites, and quantify their frameshifting probabilities. So far, detection of frameshift sites mostly relied either on conservation of the frameshift product and site and mRNA secondary structure elements^34,35^, or on ribosome profiling data^36,37^, which only detects reading frame transitions with sufficient read coverage of a given gene. Other methods, such as PRFect^38^, focus on detecting cases of PRF similar to those already known. In-vitro measurements of frameshift probabilities have primarily focused on variants of well-characterized frameshift cases or specific codon patterns (e.g. X XXY YYZ, where X, Y, Z represent nucleotides)^2,3,39^ which has limited their ability to identify novel sites. Here, we present SLIPPERRS (SLIPPery ERRor Sites), a simple mechanistic model capable of predicting frameshifting with high accuracy. Guided by the model’s predictions and using a novel cell-free assay, we identified several novel frameshift sites with probabilities exceeding those of known functional sites (Figure 1A). We analyzed 200 prokaryotic genomes *in silico* and found 18,672 putative frameshifted proteins. We found that GC-content negatively correlates with the occurrence of high frameshift sites, which tend to result in shorter and less disordered protein sequences. We detect selection to avoid the deleterious consequences of spontaneous frameshifting by selection for cryptic stop codons. Collectively, our findings demonstrate that ribosomal frameshifting is a widespread phenomenon shaping proteomes in prokaryotes with both deleterious and advantageous consequences.

**Figure 1.**
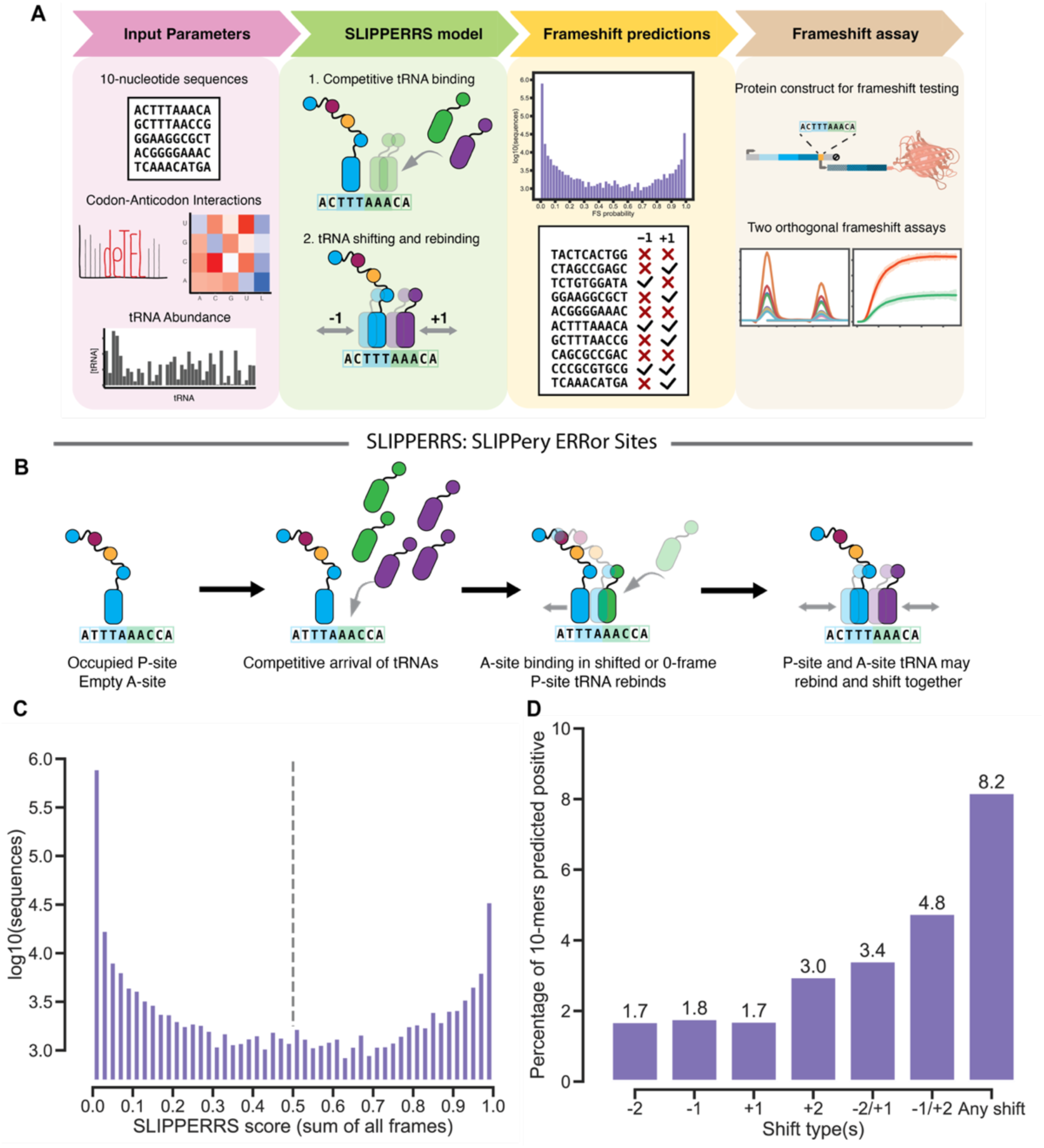
Mechanistic model of frameshifting guides identification of slippery sites. A. Computational and experimental workflow to identify slippery sites. Based on codon-anticodon interaction parameters from Landerer et al. (2024)^33^, and the cell’s tRNA abundances, the model predicts whether a 10-mer causes frameshifting. Next, 10-mers are selected for cell-free quantification of their frameshifting capability by fluorescence and mass spectrometry analysis. B. SLIPPERRS (SLIPPery ERRor Sites) is a mechanistic model predicting frameshifting by -2, -1, +1, and +2. To start, we assume that a correct (cognate) tRNA sits in the P-site, and the A-site is empty. tRNAs arrive at the ribosome in a competitive manner, with more abundant tRNAs having more chances to bind. tRNAs can bind to the A-site in the 0-frame or in a shifted frame. The latter requires the P-site tRNA to rebind and ‘match’ the shift. After a tRNA has bound to the A-site, both tRNAs may rebind and shift together (see methods). C. Histogram of SLIPPERS scores (summed over -2/-1/+1/+2 shift types) for all possible 10-mers (4^10^, excluding 10-mers containing in-frame stop codons, bin size is 0.02). Most sequences have a score close to either 0 or 1, allowing robust classification of low FS and high FS sites. The default threshold is 0.5. D. Percentage of 10-mers predicted to shift by the model (high-FS sites). Using a score threshold of 0.5, the proportion of all 10-mers predicted to cause frameshifting is between 1.5 and 3% for the four shift types, and 8.2% for any shift.

## RESULTS

### Mechanistic model of frameshifting identifies slippery sites

To systematically study frameshifting sites, we developed SLIPPERS, a simple mechanistic model of frameshifting. We model frameshifting sites of ten nucleotides (two codons and the two adjacent nucleotides on either side), which, when considering the ribosome’s P- and A-site, is the minimum sequence to model shifts by up to two nucleotides (-2, -1, +1, +2).

Our model considers two factors: Competition between tRNAs to bind to the A-site (and thus their abundance), and the binding affinity between codons (0-frame and shifted) and anticodons in P-site and A-site (Figure 1B). Our model’s *effective binding affinity* considers both the codon-anticodon sequences and the ribosome’s specific environment The effective binding affinity was obtained from a model of amino acid misincorporation (deTEL) developed previously^33^.

We model frameshifting both during decoding and during translocation. The tRNAs present in the cell compete for binding to the A-site, in either the 0-frame or a shifted frame. Binding probabilities depend on codon-anticodon affinities in both the A-site and P-site. To account for differences in the abundance of tRNAs within the cell, we scale the binding probabilities of each tRNA by its abundance. After binding, P-site and A-site tRNAs may shift and rebind together, modeling a translocation shift. Since we assume frameshifting during decoding and translocation to be independent, we sum the probabilities obtained for the two steps, and finally sum from the tRNA to the amino acid level, producing a score for each amino acid for each shift type. We do not interpret the model’s predictions as direct measures of frameshifting probabilities. Instead, we assume that higher-scoring 10-mers are more likely to function as effective frameshift sites. Consequently, the model serves as a classifier for identifying frameshift sites.

The model is general and applicable to any organisms with known relative tRNA concentrations and sufficient proteomics data to obtain parameters for codon-anticodon binding for example by using deTELpy^40^. Here, we classified all possible 10-mers (4^10^, approximately 1 million), excluding those containing in-frame stop codons, as the model is not applicable to such sequences, using *E. coli* parameters. For the majority of 10-mers, the score is close to either 0 or 1: 82% of 10-mers have a score below 0.01, 3% have a score above 0.99 (Figure 1C), making the model robust to the choice of threshold. We use a score threshold of 0.5, and consider all sequences above this threshold as predicted positive (classified as FS sites), and all others as predicted negative. With a threshold of 0.5, 1.7% - 3% of sequences (depending on the shift type) are predicted positive, and 8.2% of sequences (ca. 78,000 10- mers) are predicted positive for at least one shift type (Figure 1D).

Overall, SLIPPERS, our classifier, predicts a high number of novel frameshift sites, a subset of which we further validated experimentally.

### Method for high-throughput screening and quantification of frameshift sites

We designed an assay for the detection and quantification of frameshifting using two orthogonal readouts: relative fluorescence, and quantitative mass spectrometry (MS). The assay is based on a synthetic protein construct (Figure 2A) with three main components: the frameshift site (10-mer), two sets of peptides before and after the FS site for MS-based quantification, and the fluorescent protein mScarlet-I.

**Figure 2.**
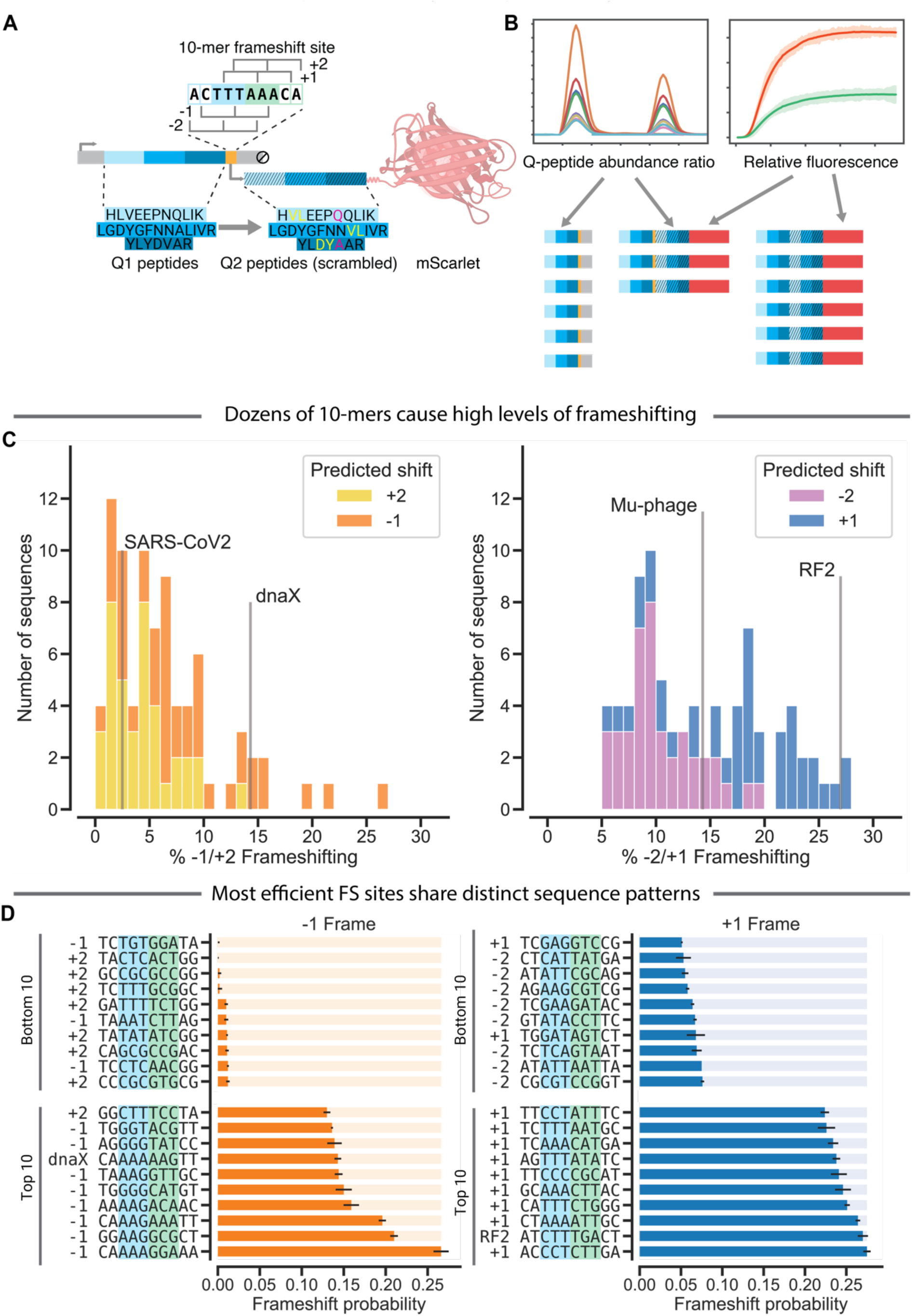
Quantitative assay discovers novel ribosomal frameshift sites that exceed efficiency of programmed frameshifting cases. A. Reporter construct design to quantify frameshifting. The 10-nucleotide sequence is inserted (surrounded by spacers) between the coding sequences of two sets of peptides (Q1 and Q2), made up of three pairs of highly similar peptides. The second set of peptides is encoded in the shifted frame (-1/+1), followed by mScarlet-I. In the 0-frame as well as the third, non-tested frame, the 10-mer is followed shortly by stop codons. Hence, mScarlet-I and Q2 peptides are only produced upon frameshifting, allowing the quantification of frameshifting probabilities. B. Frameshifting is quantified by two orthogonal methods. Translation of the protein construct’s coding sequence produces two products: The shorter non-FS product (Q1, 10-mer, spacers), and the large FS product (Q1, 10-mer, spacers, Q2, mScarlet-I). The ratio of the total amounts of Q1 and Q2 peptide counts measured by mass spectrometry is a direct readout of the FS probability. The fluorescent intensity relative to a control sequence expressing mScarlet-I independent of frameshifting provides a second estimate of FS probability. C. Histogram of in-vitro FS probabilities measured in cell-free expression system quantified by relative fluorescence (-1 frame (left) and +1 frame (right)). Many tested 10-mers exhibit >10% frameshift probability, and some cause higher -1 or +1 frameshifting than the well-known programmed ribosomal frameshifting (PRF) sites of dnaX and RF2, respectively. D. In-vitro FS probabilities of the 10 most and 10 least efficient FS sites in -1 (left) and +1 (right) frame. Black error bars indicate standard deviation of three technical replicates. The shift type predicted by SLIPPERRS is indicated next to the sequence, except for PRF sequences. P-site is highlighted in cyan, A-site in green. Most efficient FS sites show sequence similarities, particularly in the P-site codons: of the efficient -1 sites, the P-site codons and the 5’ nucleotide are extremely purine-rich, while of efficient +1 sites, many P-site codons and the 3’ nucleotides form a tetramer that allows +1 rebinding with only a wobble mismatch.

We designed a fluorescent reporter assay, that is based on an out-of-frame-encoded mScarlet-I which is only produced upon frameshifting. Since the amount of mScarlet-I is proportional to the fluorescence intensity (Figure S3), we can quantify frameshifting probabilities by comparing the fluorescence intensities to a non-shifting control sequence, where all components are encoded in the 0-frame and mScarlet-I is constitutively expressed (‘0-control’). The ratio between the endpoint fluorescence values in the 0-control and a given test sequence gives the estimated frameshifting probability (Figure 2B, Figure S2).

To verify that the relative fluorescence values genuinely reflect frameshift probabilities, we used an orthogonal frameshift quantification using mass spectrometry (MS). We designed three pairs of ‘quantitative peptides’ (Q-peptides), based on previous work by the Shevchenko lab^41,42^. For each pair, one peptide is positioned in the 0-frame, before the FS site (‘Q1’), and the other one after the FS site, in the shifted frame (‘Q2’). The two peptides of each pair can be distinguished by MS, but are similar enough that their signal intensities can be directly compared, allowing relative quantification. By quantifying all Q-peptides by MS, we can calculate the ratio of abundance between Q2 and Q1 for each pair (Figure 2B). The frameshifting probability is then calculated as the median of the three individual Q2/Q1 peptide ratios. Frameshift probabilities quantified by MS and by fluorescence correlate well (Pearson’s *R* = 0.73, Figure S4), validating our high throughput fluorescence assay and suggesting that it is not affected by different expression levels of individual reporters.

While our fluorescence reporter assay allows estimation of FS probabilities with high throughput and sufficient dynamic range (Figure S3), it may suffer from non-canonical translation events. We wanted to control for potential frameshifting and alternative initiation^43^. To correct for any frameshifting occurring on the 0-control sequence, we included two additional constructs containing the same 10-mer as 0-control, but with Q2 and mScarlet-I in the -1 and +1 frame, respectively. We then use the total fluorescence from all three frames as the new reference to calculate relative fluorescence and thus frameshift probabilities.

Alternative initiation sites, before or at the beginning of mScarlet-I, would lead to an increase in the amount of mScarlet-I (and fluorescence) independently of frameshifting. Indeed, comparing the absolute amounts of Q1 peptides produced to the relative fluorescence, we detect excess fluorescence (see Methods for details) indicative of fluorescence independent of frameshifting, which we assume is due to alternative initiation. In our assay, alternative initiation accounted for 7.2 percentage points of relative fluorescence (Figure S5, Figure S6), which we then subtract from the measured relative fluorescence to calculate frameshift probabilities.

Given that measured frameshifting probabilities are likely influenced by factors such as temperature or co-factor concentrations, we interpret our measurements as relative, focusing on the comparative ability of different 10-mers to induce frameshifting. This assay enables high-throughput identification of effective frameshift sites.

### Discovery of novel slippery sites surpassing known programmed frameshift rates

In order to experimentally test SLIPPERRS’ predictions and validate novel FS sites, we designed a 10-mer sequence library. We randomly selected 160 sequences: 30 positive predictions each for -2, -1, +1, and +2 shifting, and 10 negative predictions for each shift type. We excluded trivial FS sites (e.g. the XXXYYYZ pattern or repetitive 10-mers) and selected non-redundant sequences only (< 80% identity, i.e. > 2 nt differences). In addition, we included a few known frameshift sequences as controls including *E. coli* RF2 (+1)^25^, *dnaX* (-1)^27^, the Mu-phage (-2)^44^, SARS-CoV2 (-1)^45^ (see methods for details), as well as a 10x thymine repeat as a presumably very slippery site.

In total, we tested 165 10-mers for frameshifting using our assay, expressing the reporter construct in the prokaryotic cell-free system PURExpress. We chose a cell-free system to increase sensitivity of detecting frameshift events. PURExpress is composed of purified ribosomes, tRNAs, and translation factors, providing a highly controlled environment for studying translation mechanisms without cellular complexity. Most importantly, it lacks endogenous proteases and protein degradation machinery that would likely eliminate frameshifted protein products. We can thus directly study the ribosome’s error rate.

The majority of sequences in the library caused detectable levels of frameshifting, with FS probabilities of up to 28% (Figure 2C). The known FS sites of *dnaX*, RF2, and phage Mu were all among the most efficient FS sites in their respective frames, with 14%, 27%, and 14% frameshifting, respectively. The SARS-CoV2 sequence, which naturally works in a eukaryotic system, gave only 2.5%. Interestingly, several sequences were even more efficient than known functional FS sites: Six sequences shift into the -1 frame more efficiently than the *dnaX* site; the FS probability of 36 -2/+1 sites exceed that of the Mu-phage site; and one -2/+1 site has a higher FS probability than the RF2 site. As expected^11,46,47^, we see a clear trend of -2/+2-shifts having a lower probability than -1 and +1. We note that the annotation of +2 and -2 sites is based on the model predictions, since our reporters cannot strictly distinguish -2 from +1 and -1 from +2 shifts. Given the trends we observe, we assume that the majority of frameshifting we observe is due to -1 and +1 shifting, and we will refer to the sequences in the -2/+1 frame and -1/+2 frame as -1 and +1 sequences hereafter.

We see clear sequence differences between the least and most efficient FS sites, as well as between the most efficient -1 and +1 sequences (Figure 2D). In the most efficient -1 sites, the P-site and the preceding nucleotide are rich in purines (G and A), and the eight most efficient -1 FS sites have one of only three P-site codons: AAA, GGG, and AAG. In contrast, the most efficient +1 sites have a high proportion of A and T. In addition, the P-site and the 1st nucleotide of the A-site often form a tetramer that would allow +1 rebinding with only one mismatch, such as CTT_T, AAA_A, or TTT_C (underscore delineates codons). These observations indicate that the P-site is a deciding factor in frameshifting. In summary, we present one of the largest systematic quantifications of frameshift probabilities, and identify highly efficient novel frameshift sites, some exceeding the frameshift probabilities of known programmed sites.

### Dual frameshift mechanism observed at the SARS-CoV2 frameshift site

While the reporters (fluorescent and MS-based) assays measure FS rates, peptides that cover the FS site can reveal the molecular mechanism of frameshifting, specifically the direction of the shift (-2 vs. +1 / +2 vs. -1), as well as whether the FS occurred at the translocation or the decoding phase. For the SARS-CoV2 10-mer, we identified two distinct peptides covering the FS site using mass spectrometry (Figure 3). While both of them indicate a -1 shift at the 10- mer, one matches a shift from 0 to -1 one codon earlier than the other. While we cannot exclude a shift by the same mechanism at two neighboring codons, we surmise that the different sequences are due to shifts at the same position, but during the translocation and decoding phase of elongation, respectively. While dual mechanisms have been described in analogous frameshift systems^11^, our work provides direct evidence of both decoding and translocation frameshift mechanisms. Notably, both mechanisms are implemented in SLIPPERRS, giving support to our mechanistic model.

**Figure 3.**
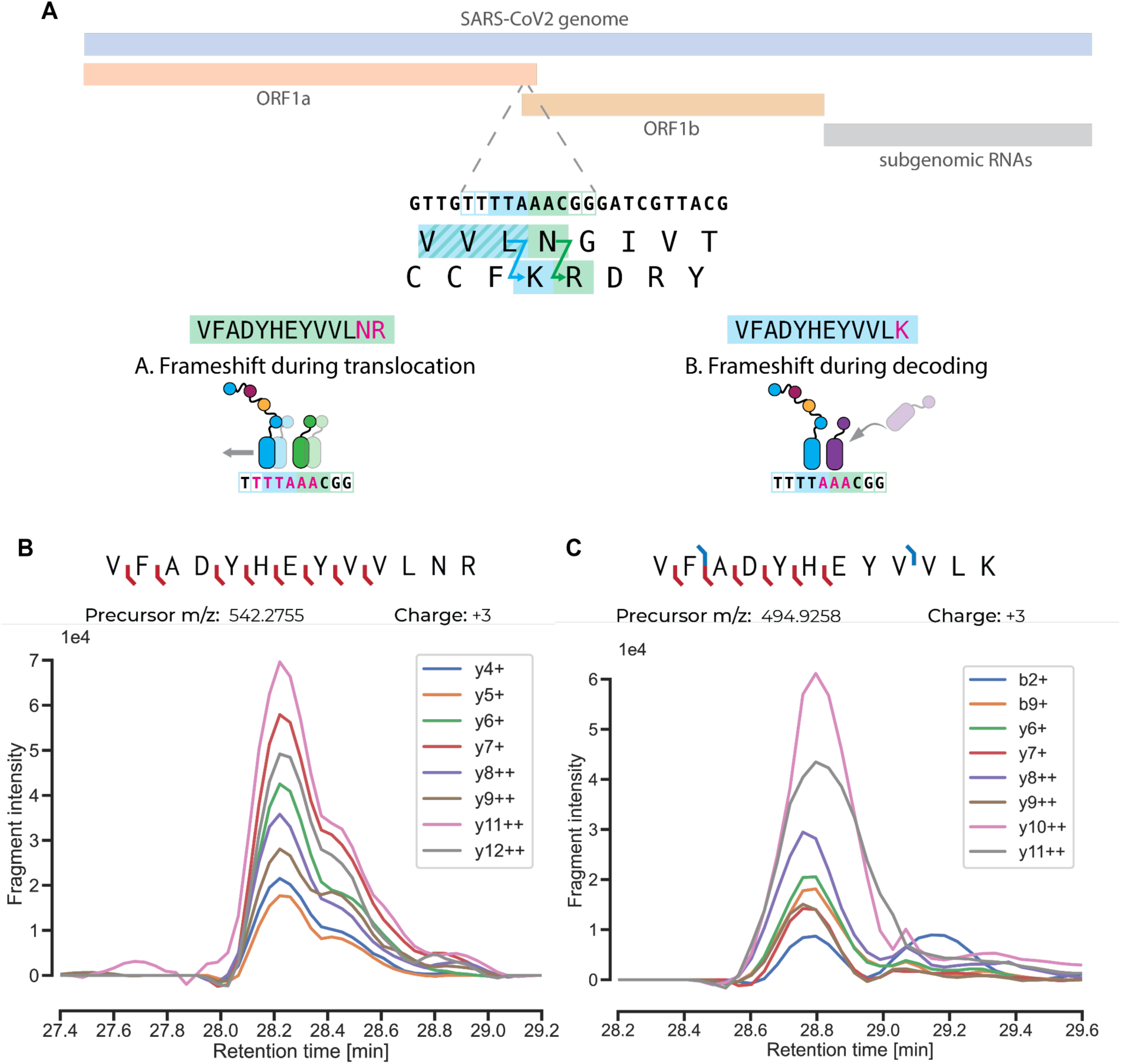
Evidence for dual frameshift mechanism – via translocation and decoding – at a single site. A. The 10-mer taken from the SARS-CoV-2 genome as well as the sequence context in the frameshift quantification construct is highlighted: TTTTAAACGG. Targeted MS detected two peptides originating from a frameshift at the 10-mer, VFADYHEYVVLNR and VFADYHEYVVLK. The LNR peptide matches a shift during translocation, while the LK peptide corresponds to a shift during decoding, at the exact same site in the mRNA. B. Mass spectrometry evidence for the peptide VFADYHEYVVLNR indicating a translocation shift. The peptide was detected by targeted acquisition. Fragments with masses matching the peptide’s y-ions from y4 to y12 (with the exception of y10) were detected. The ions’ chromatographic peaks are highly congruent, indicating they originate from the proposed peptide. C. Mass spectrometry evidence for the peptide VFADYHEYVVLK indicating a decoding shift. The peptide was detected by targeted acquisition. Fragments matching the peptide’s y-ions from y6 to y11 as well as the b2 and b9 ions eluted at the same retention time, confirming the presence of the peptide. Peptide annotations generated with Interactive Peptide Spectral Annotator^86^.

### SLIPPERRS classifies frameshifting sequences with high accuracy

Using the experimentally characterized FS sites in our library, we could test the accuracy of SLIPPERRS. Our library contains 82 sequences in the -1 frame (-1 and +2 shifts) and 82 sequences in the +1 frame (-2 and +1 shifts). We only used predictions for -1 and +1 shifting, and we did not separately predict -2 and +2 shifting, for two reasons: First, our assay does not identify the shift type, only the reading frame. Second, the model performs significantly better when we only use the -1 and +1 scores (Figure S7).

With a default score threshold of 0.5, SLIPPERRS classifies 13 sequences in the -1 frame, and 14 sequences in the +1 frame, as frameshifting. For both frames, the predicted positive sequences have higher experimentally determined frameshift probabilities than the negative predictions (p = 3.8e-6 for -1; p = 3.8e-8 for +1, two-sided independent t-test; Figure 4A), confirming that the model can distinguish sequences with high frameshift probabilities.

**Figure 4.**
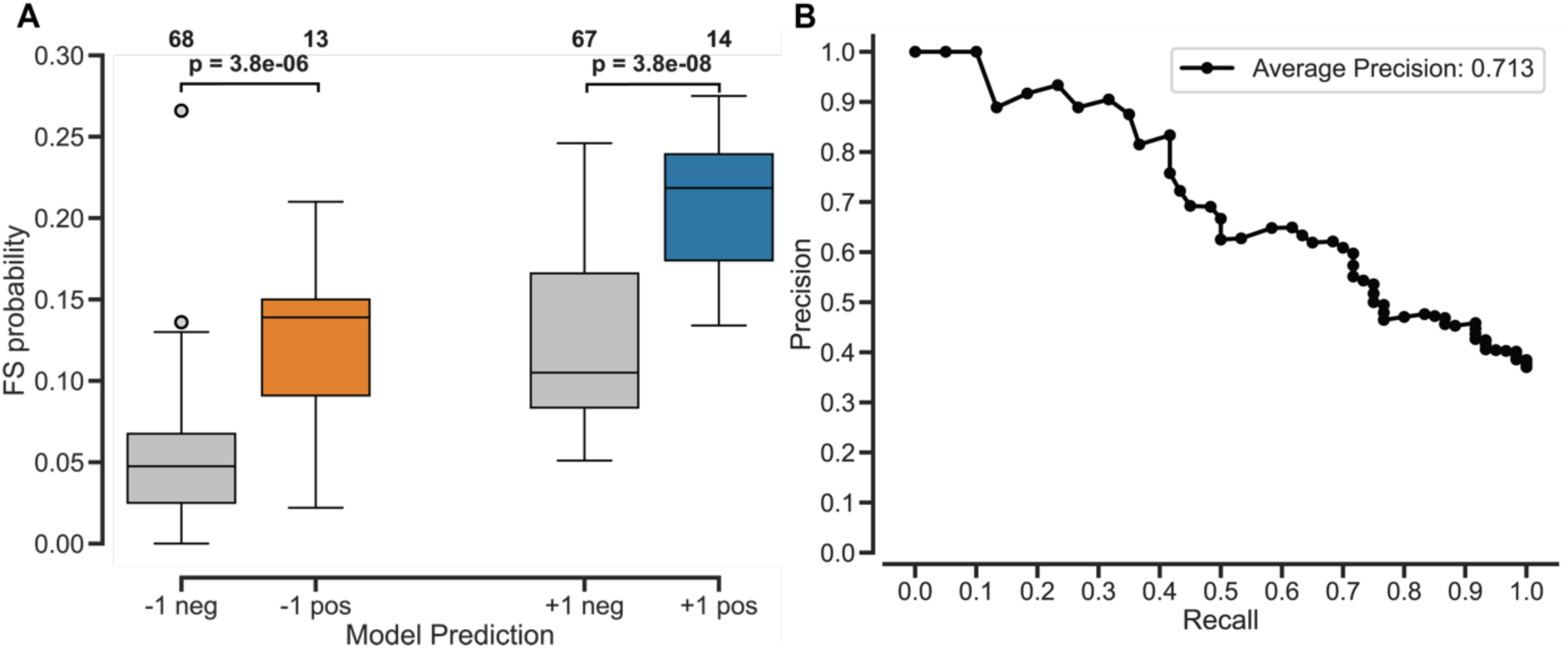
SLIPPERRS classifies frameshifting sequences with high accuracy. A. Predicted FS sites cause higher frameshift probabilities *in vitro* than negative predictions. Experimentally measured 10-mers are grouped according to their frame and the model’s classification. P-values were determined by two-sided independent t-test, group sizes indicated above. FS sites predicted by SLIPPERRS have significantly higher frameshift probabilities than negative predictions. B. Precision-Recall curve of model predictions for the experimentally tested library. Based on the frameshifting probabilities measured by fluorescence, 10-mers were classified as ‘high-FS’ (positive) and ‘low-FS’ (negative) sites (<10.4%). SLIPPERRS achieved an average precision across all score thresholds of 0.713 when predicting high-FS 10-mers and, at the default threshold of 0.5, an overall accuracy of 73%.

We next classified the library sequences based on the experimentally measured frameshift probabilities into ‘high-FS’ and ‘low-FS’ site categories. We selected a threshold (10.4%) based on the shape of distributions of both -1 and +1 FS probabilities, to select the tail of the distributions and to include all prokaryotic PRF sequences (Mu-phage, RF2, and *dnaX*). As a result, 12 out of 82 sequences in the -1 frame and 49 out of 82 sequences in the +1 frame are classified as ‘high-FS’. This classification allowed us to test the model’s performance more quantitatively: SLIPPERRS is able to predict high-FS sites with an average precision of 0.71 (Figure 4B). At the optimal score threshold of 0.27, SLIPPERRS achieves an F1 score of 0.55, and, at the default threshold of 0.5, a slightly lower F1 score of 0.5 and an accuracy of 73%. We thus show that our mechanistic model, despite not being trained on frameshift data in any way, is able to predict efficient FS sites with high accuracy.

### Codon patterns and slippery P-site are predictive of frameshift probabilities

To gain a deeper understanding of the factors influencing FS probabilities, we sought to identify sequence traits predictive of high frameshifting probabilities as measured by our experimental assays. We identified three codon patterns that are strongly predictive of frameshifting probability (Figure 5B):

1. Sequences with an XXX codon in the P-site have much higher FS probability (codons with the highest average frameshifting probability were AAA, UUU, and GGG, Figure 5A).
2. An XYX codon in the P-site is associated with lower FS probability (e.g. GCG, UCU, and CGC, Figure 5A).
3. Sequences with an XUU codon have higher average FS probability (Figure 5A).

**Figure 5.**
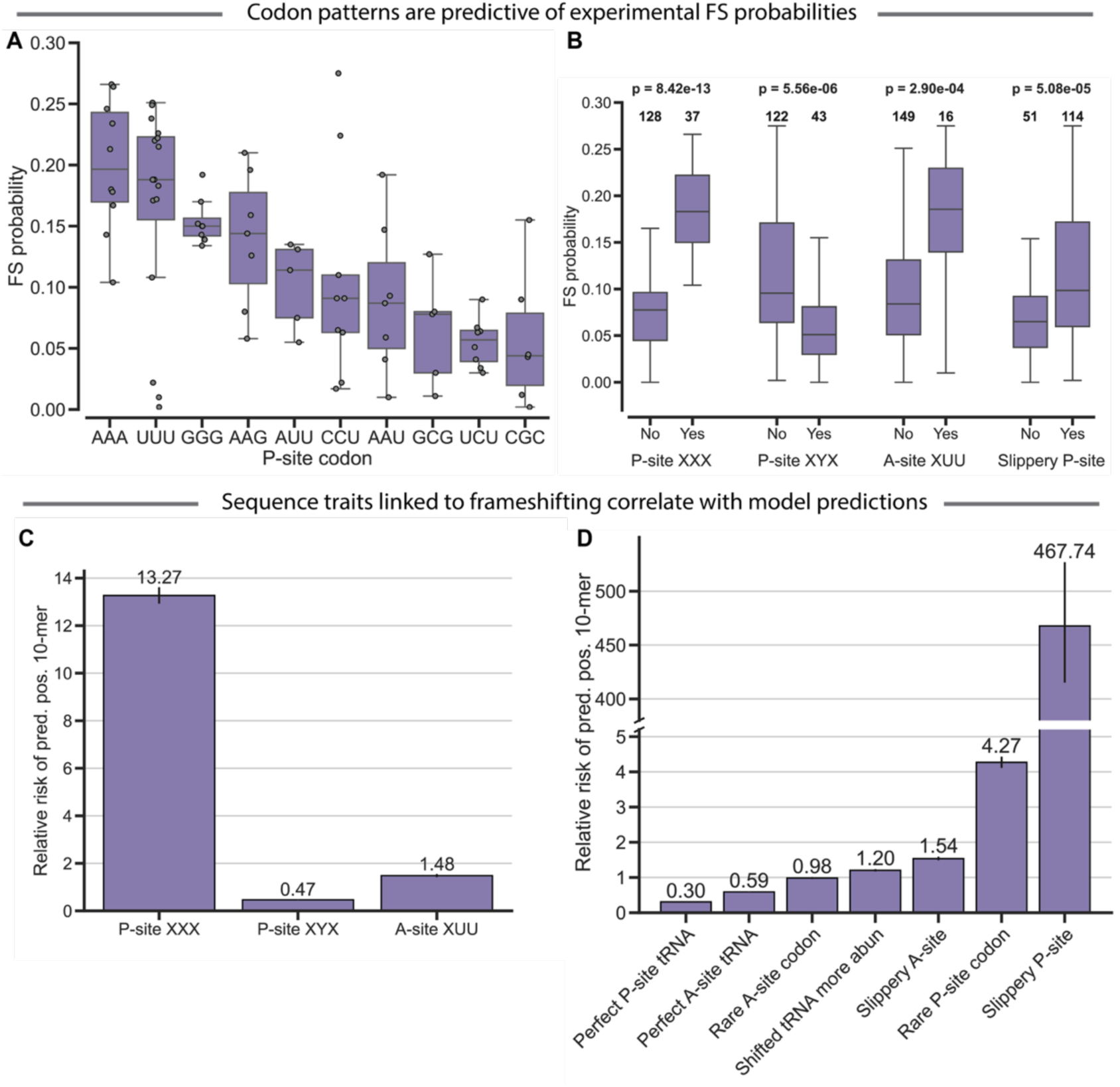
Codon patterns and slippery P-site are predictive of frameshift probabilities. A. P-site codons correlate with measured frameshift probability. Experimentally tested 10-mers are grouped by their P-site codon. Only groups with at least five sequences are shown. The codon groups are sorted by their median frameshift probability. 10-mers with repetitive codons (AAA, UUU, GGG) show higher than average FS, while GCG, UCU, and CGC are associated with low FS. B. Codon patterns are predictive of frameshift probability. 10-mers are grouped depending on whether their P-site or A-site codon match one of four patterns. P-values were determined using two-sided independent t-test. An XXX codon in the P-site or an XUU codon in the A-site are associated with higher FS probabilities, an XYX P-site codon with lower FS probabilities. A ‘slippery’ P-site (facilitating rebinding of the P-site tRNA, see Methods for details) is associated with higher FS probabilities, as well. C. Codon patterns predictive of experimental FS also correlate with model predictions. Bars indicate the relative risk (change in proportion of 10-mers that are predicted to frameshift), i.e., a relative risk of 2 would indicate a 2-times higher likelihood of being predicted positive if the 10-mer has a certain property, while 1 would indicate no influence of the property. Black lines indicate the 95% confidence interval (see Methods for details). The three codon patterns identified from experimental data influence model predictions in a similar way, with P-site XXX and A-site XUU increasing FS and P-site XYX decreasing it. D. Frameshift-promoting sequence properties correlate with SLIPPERRS predictions. 10-mers were tested for properties associated with frameshifting, such as usage of rare codons or the ability of the 10-mer to allow tRNA rebinding (see Methods for details). Bars indicate the relative risk, i.e. change in proportion of 10-mers that are predicted to frameshift between sequences with vs. without the property. A ‘slippery’ A-site or P-site (such as X_XXY), a rare P-site codon, and the tRNA cognate to a shifted codon being more abundant than the 0-frame tRNA promote frameshift prediction.

These patterns fit with our intuition about the mechanism of frameshifting: We expect that frameshifting requires a rebinding of the P-site tRNA. A codon of the pattern XXX would make this more likely, since a (perfectly matching) anticodon would have at most a single mismatch after shifting by one nucleotide. In contrast, an XYX codon in the P-site would mean that a shift by one nucleotide, again assuming a perfect match between codon and anticodon, results in at least two mismatches. In contrast, codon patterns in the A-site seem to be less important (Figure S8). These observations point to a ‘slippery’ P-site as the most important determinant of FS.

Consequently, we classified the P-sites of all our 10-mers as slippery or not, where slippery means that a tRNA cognate to the 0-frame might also bind to the shifted frame (see Methods for details). We clearly see higher FS probabilities for 10-mers with slippery P-sites (Figure 5B). This trend holds for the A-site, although it is weaker. (Figure S9). This agrees with our earlier observation that we can see very clear patterns for the P-sites of the most efficient 10- mers, but not necessarily for their A-sites (Figure 2D). Other traits such as a perfectly matching tRNA, or the relative synonymous codon usage (RSCU), seem to be less important (Figure S9). In summary, we identify sequence properties predictive of prokaryotic frameshifting that support our modeling assumptions and give insight into the mechanism of frameshifting.

### SLIPPERRS predictions recapitulate frameshift-promoting sequence traits

Following from the analysis of the sequence traits that influence the experimentally tested frameshift probabilities, we investigated whether similar correlates could be identified for the SLIPPERRS scores of -1 and +1 shifting. We used relative risk as a measure of the influence of a sequence trait, where relative risk is the ratio of the proportion of 10-mers predicted positive (classified as FS sites) with and without the trait, with 1 indicating no effect (see Methods for details).

The three codon patterns predictive of *in vitro* FS probabilities likewise influence the model predictions: For P-site XXX and A-site XUU, the relative risk is clearly above 1, indicating that the model is more likely to predict sequences with these codon patterns as positive. For P-site XYX, we see the opposite effect (Figure 5C).

Building on this and inspired by a similar analysis^4^, we investigated more complex sequence traits that we assume to be connected to frameshifting (Figure 5D, see Methods for details). Among these traits, a slippery P-site has by far the largest influence on the model’s predictions. In addition, a slippery A-site, a rare P-site codon and an abundant tRNA decoding the shifted frame make 10-mers more likely to be predicted positive. In contrast, a perfect match between codon and anticodon in either the P-site or the A-site lead to fewer classifications as FS sites (Figure 5D). We note that none of these sequence traits are explicitly part of SLIPPERRS.

However, it seems they are implicitly reflected in the way we model frameshifting, and the model’s behavior reflects the state of knowledge about ribosomal frameshifting.

### High-frameshift sites are widespread and negatively correlate with GC content in prokaryotic genomes

Our dataset of verified frameshifting sites allowed us to assess the extent of frameshifting in prokaryotic genomes; specifically, where frameshift sites occur, and what their consequences are. Since we quantified frameshift probabilities in a cell-free system containing *E. coli* translation machinery, we selected 200 *Gammaproteobacteria* species including *E. coli*.

We scanned these 200 genomes for the experimentally characterized 10-mer sites, classified as either high-FS sites (N=60) or low-FS sites (N=102). We only included occurrences of the 10-mers within coding sequences, and within frame, such that what we consider the 10-mer’s P-site and A-site codon are in the gene’s 0-frame. We identified 18,672 occurrences of high-FS 10-mers and 32,403 occurrences of low-FS 10-mers in total amongst our *Gammaproteobacteria* dataset. Approximately 2.4% of all genes (18,306 / 770,151) contain a high-FS site (Figure 6A).

**Figure 6.**
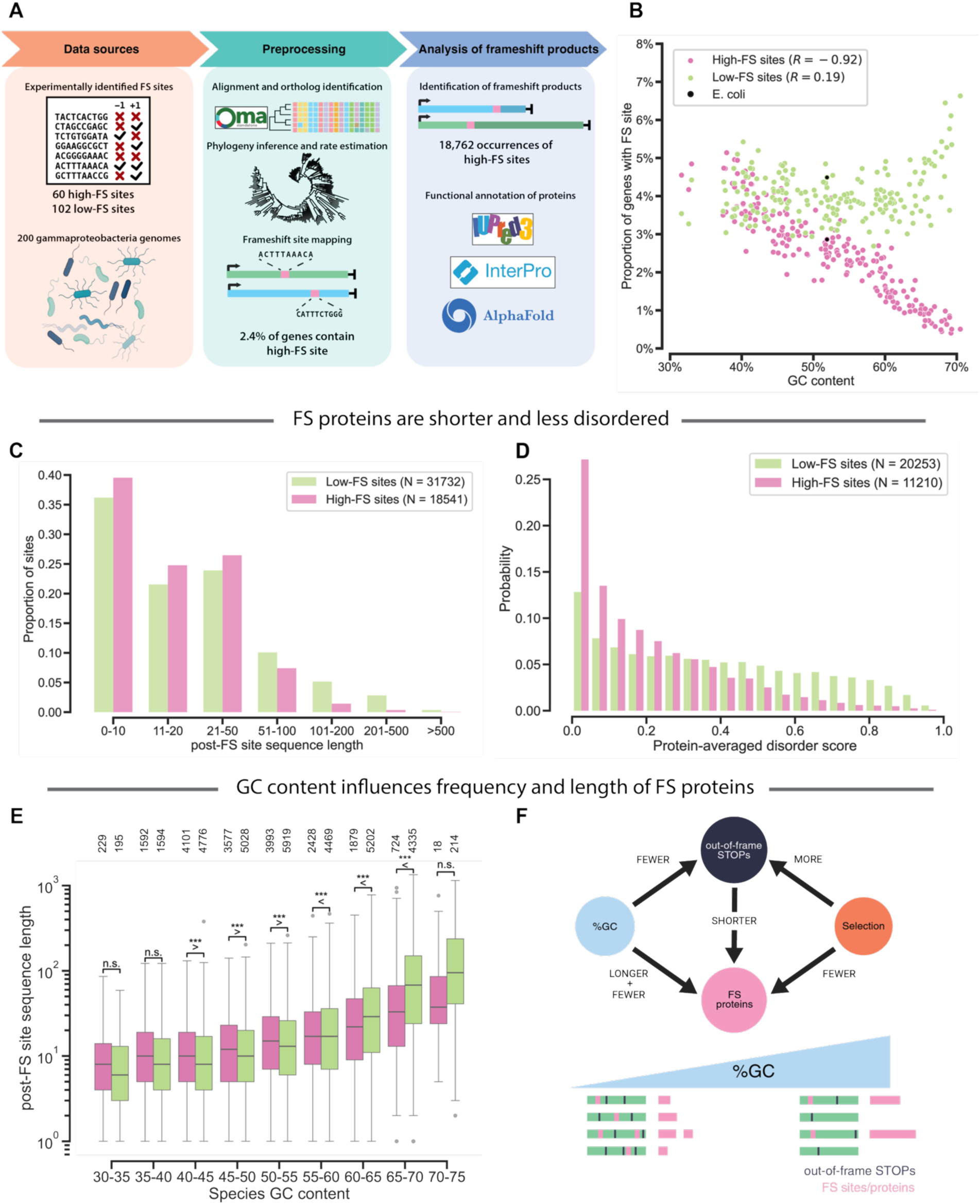
Evidence of selection against the consequences of frameshifting in *Gammaproteobacteria*. A. Schematic overview of the analysis of frameshift sites in prokaryotic genomes. FS sites (10-mers) from the *in vitro* library are mapped to the coding sequences of 200 *Gammaproteobacteria* species. Coding sequences are aligned, orthologs identified and a phylogeny inferred. The positions and occurrence patterns of FS sites, as well as the hypothetical frameshift protein products, are analyzed and functionally annotated. B. Scatterplot of the proportion of genes with high-FS and low-FS sites (per species) versus the GC content. The proportion of genes with high-FS sites is strongly negatively correlated with the GC content, while for low-FS sites, the correlation is weak (*R:* Pearson correlation coefficient). C. Histogram of the distribution of the length of the protein sequence past the FS site for all hypothetical FS products in *Gammaproteobacteria,* excluding cases where the FS would fuse two genes, for low-FS (N = 31,732) and high-FS (N = 18,543) sites. Products of high-FS sites are shorter on average compared to products of low-FS sites (two-sample, two-sided Kolmogorov-Smirnov test: p = 3e-86). In particular, long FS products (>100 residues past the FS site) are very rare for high-FS sites. D. Histogram of protein-averaged disorder scores of post-FS-site regions of hypothetical FS products in *Gammaproteobacteria*. For the predicted protein products of all FS sites detected in the *Gammaproteobacteria* dataset, disorder was predicted using IUPred2A-long, for low-FS (N = 20,253) and high-FS (11,212) sites. Only the sequence region after the FS site was considered, and only sequences where this region was at least 10 residues long, excluding fusions. Protein products of high-FS sites are much less disordered than products of low-FS sites (two-sample two-sided Kolmogorov-Smirnov test: p ≍ 0). E. Histogram of post-FS site sequence length of frameshift proteins, binned by the GC content of the species. The protein length increases quickly at GC contents of 60% or more. In the high-GC range, where proteins tend to be longest, products of high-FS sites are significantly shorter than those of low-FS sites. P-values from two-sided independent t-test are indicated: ***: < 0.001, **: < 0.01, *: <= 0.05, n.s.: > 0.05. F. Scheme of the influence of GC content, occurrence of out-of-frame stop codons and selection on the frequency and length of frameshift proteins. A high GC content leads to a lower frequency of cryptic stop codons and effective frameshift sites (both of which are TA-rich). The lower frequency of stop codons leads to (mostly deleterious) longer frameshift products. This in turn suggests that in organisms with high GC content, there is stronger selection for additional cryptic stop codons.

As described above, high-FS sites in our dataset have a higher proportion of T/A. This composition bias means that, simply by chance, they should be less likely to occur in genomes with a high GC content. Indeed, we see a strong negative correlation between a species’ GC content and the proportion of genes containing a high-FS site (Pearson’s R = -0.92), while there is only a weak relationship for the low-FS sites (Figure 6B). The proportion of genes in a given species that contain such a site ranges from below 1% to ca. 5%, indicating that efficient FS sites are frequent in these genomes.

### Products of high-frameshift sites are shorter and less disordered

We next investigated the hypothetical proteins that would be produced by a frameshift at these sites. These proteins would miss a part of the canonical protein sequence, replaced by a novel sequence region encoded by the shifted frame up to the next shifted-frame stop codon. We saw significant differences between the hypothetical proteins produced by high-FS and low-FS sites: On average, efficient FS sites produce shorter frameshifted proteins, and in very few cases would the protein be extended by more than 200 amino acids. In contrast, about 3% of low-FS sites would lead to a frameshifted sequence between 200 and 500 residues, in the range of an average full-length prokaryotic protein (Figure 6C). The most striking difference between efficient and inefficient sites was revealed by analyzing the disorder content of the frameshift proteins: Proteins produced by high-FS sites would be much less disordered (as predicted by IUPred2A, see methods): The average disorder scores (per protein) for the products of high-FS sites are much lower than for low-FS sites (Figure 6D). This trend holds for per-residue scores and protein disorder content (Figure S10). Globally, the most efficient frameshift sites would produce shorter and more ordered proteins than expected. Since larger proteins are energetically costly to synthesize and disordered proteins are more susceptible to toxicity, e.g. via aggregation, it would be beneficial for organisms to avoid the random production of large, disordered proteins.

### Selection for cryptic stop codons mitigates longer frameshift products in GC-rich bacteria

We saw before that due to their high TA content, high-FS sites occur less often in genomes with higher GC content (Figure 6B). Similarly, with higher GC content we expect fewer ‘cryptic’ (out-of-frame) stop codons. Since the occurrence of out-of-frame stops determines the length of frameshifted proteins, this prompted us to investigate the relationship between GC content and FS protein length. As expected, the length of frameshift proteins increases with GC content, although this trend only becomes pronounced at high GC content of 60% or more. However, in this GC range, where most of the very long frameshift products occur, the products of low-FS sites are significantly longer than those of high-FS sites (Figure 6E). It is possible that species with high GC content, where FS sites would be rarer, but their products much larger, have been subject to selection to increase the frequency of out-of-frame stop codons (Figure 6F).

Together, these observations suggest that GC content and out-of-frame stop codons are under selection pressure to reduce the production of longer, potentially deleterious proteins resulting from frameshifting.

### Naturally occurring frameshift sites cut off domains, fuse canonical genes, and create large, structured extensions

We next investigated how frameshifts affect the structure and function of the proteins. The consequence of a frameshift can be categorized according to how it affects the protein’s functional domains (Figure 7A): Depending on where the frameshift happens and what is encoded by the frameshifted region, the FS product might retain all, at least one, or none of the canonical protein’s domains. We refer to these scenarios as ‘preserved’, ‘truncated’, and ‘destroyed’, respectively. In addition, the frameshift region might contain a new domain not originally present (‘FS domain’). Finally, a frameshift might lead to a fusion with a gene adjacent in the genome (‘fused’).

**Figure 7:**
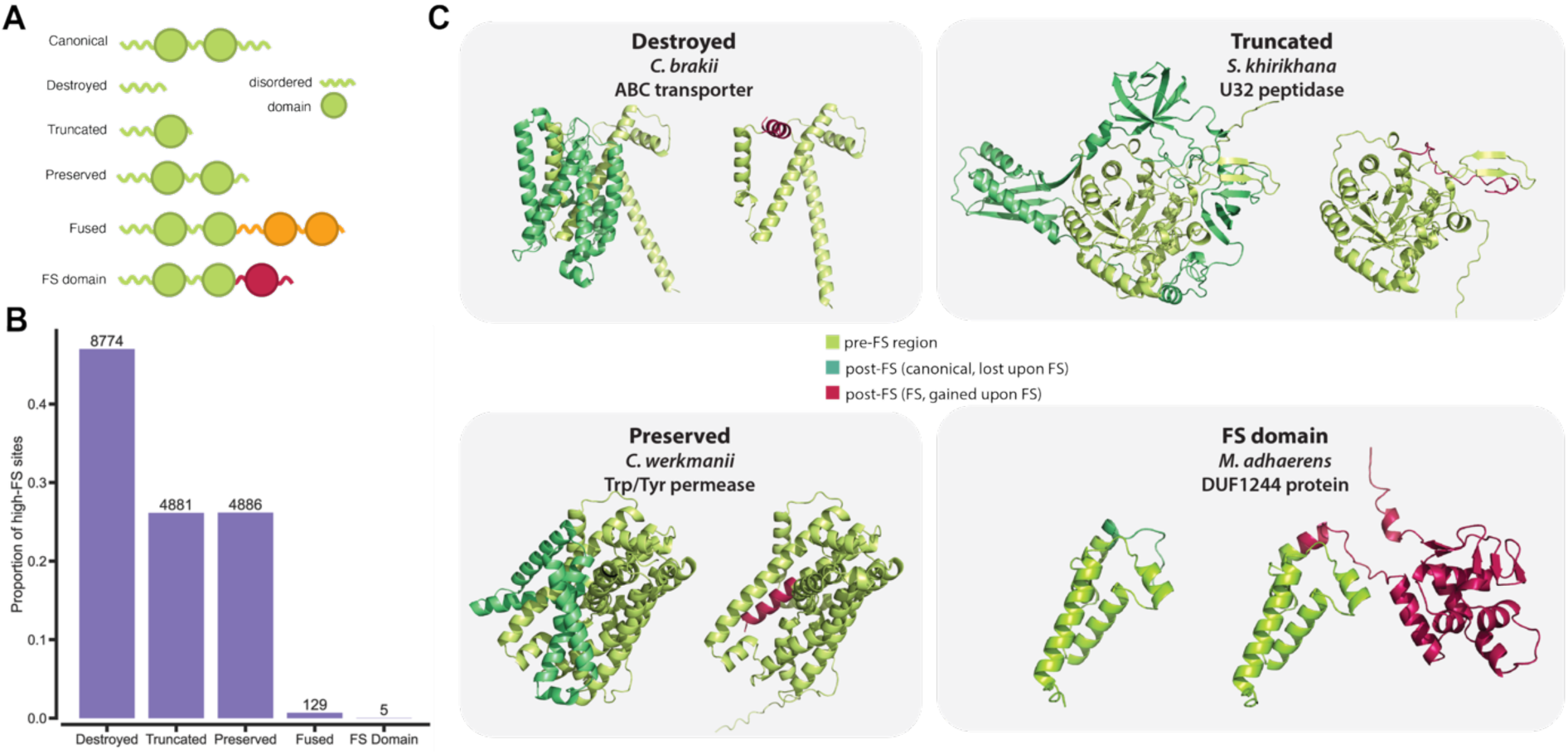
Frameshift sites diversify the proteome. A. Scheme of possible consequences of frameshifting on the protein. The protein is conceptualized as a sequence of ordered domain and disordered regions. The canonical protein might be destroyed, truncated or preserved, depending on whether all, some, or none of its domains are lost upon frameshifting, respectively. In addition, the frameshift might lead to a fusion with another gene, or the FS region might give rise to a new domain of its own. B. Proportions of categories of frameshift consequences in *Gammaproteobacteria*. Displayed are the proportions of the predicted products of high-FS sites that fall into each of the categories. Categories were assigned based on Pfam domains in the canonical and frameshifted proteins as detected by Interproscan. Apart from fusions with another gene, cases where the frameshifted region contains a domain are extremely rare (N = 5). C. Displayed are the Alphafold3 predictions of 3D structures for four *Gammaproteobacteria* proteins containing a high-FS site. The regions pre-FS site (unaffected by FS) and post-FS for the canonical protein (lost upon FS) and the FS product (gained upon FS) are marked in green, blue-green, and red, respectively. *Upper left:* ABC transporter WP.016156150.1 from *C. brakii*, with a +1 FS site at codon 130, leading to the destruction of the only domain. *Upper right:* U32 peptidase WP.126165837.1 from *S. khirikhana* with a +1 FS at codon 315, truncating the protein. *Lower left:* WP.137347863.1 from *C. werkmanii*, with a +1 FS at codon 318, preserving the domain. *Lower right:* DUF1244-containing protein WP_014576245.1 from *M. adhaerens* with a -1 FS at codon 80, adding a new domain to the protein.

Using Interproscan to annotate functional domains, we classified the hypothetical frameshift products of all high-FS sites (N = 18,672) in *Gammaproteobacteria* into these categories. The most common case is the loss of all functional domains (‘destroyed’, ca. 47% of cases), followed by preserved and truncated (both 26%) canonical proteins (Figure 7B). In 129 (0.7%) cases, the frameshift leads to a fusion with another gene. Cases where the frameshift produces a domain not present in the canonical protein are extremely rare: Excluding fusions, only five frameshift proteins contain a new domain (see Figure 7C for examples).

Using *Gammaproteobacteria* as an example, we thus show how frameshifting can contribute to proteome diversity.

## DISCUSSION

To date, investigation of ribosomal frameshifting sites has largely focused on known programmed frameshift sites and variants thereof^2,3,39^. Here, we systematically explored frameshift sites across organisms to study their impact on sequence evolution and proteome diversification. We present SLIPPERRS, a computational model, and an experimental approach to quantify frameshifting probabilities high-throughput allowing proteome-wide assessment of the impact of frameshifting. We measured in-vitro frameshift probabilities of over 160 10-mers. We can assume that many of the 10-mers we investigated are genuine effective frameshift sites, since they cause comparable or higher frameshifting than functional prokaryotic sites: *dnaX*, Mu-phage, and RF2. In addition, our FS probabilities for these sites agree with previous studies: The *dnaX* site on its own, without stimulators, previously caused 16% FS *in vitro*^11^, compared to 14% in our assay. For RF2, previous measurements, which included the 5’ stimulatory sequence, are 33% *in vivo*^48^ and 50% *in vitro*^24^, while we, using only the 10-mer, detect 27%. The Mu-phage site caused FS levels very similar to *dnaX,* and to our knowledge, this is the first FS quantification of that site.

Despite these agreements, the frameshift probabilities we measured in our cell-free system are higher than expected. This is likely because we measure frameshifting in a bottom-up cell-free expression system. Indeed, we tested a subset of our sequences *in vivo*, by transforming bacteria with the same frameshift testing constructs. While the observed FS probabilities *in vivo* and *in vitro* correlated, *in vivo* probabilities were significantly lower (Figure S11). These differences in the fidelity of translation between *in vivo* and *in vitro* studies have been observed before^13^. It is possible that the PURE system lacks proteins or other components that reduce translation errors in general or frameshifting in particular *in vivo*. Candidates for such proteins are the elongation factor 4^49^, the elongation factor P^50^, or *smrB*^51^. Despite the differences in absolute values, the correlation between *vivo* and *vitro* measurements (Figure S11) as well as the validation by mass spectrometry (Figure S4) make us confident that our assay accurately measures the relative ability of sequences to cause frameshifting and thus identifies genuine efficient frameshift sites.

The large-scale identification of high- and low-FS sites allowed the identification of sequence properties that are predictive of frameshifting. Some of these agree with previous knowledge and intuition about the frameshift mechanism, such as the high number of repetitive (XXX) P-site codons among the most efficient sites. Others were less expected, such as the frameshift promoting effect of A-site XUU, or the high efficiency of -1 sites not matching the classic XXXYYYZ pattern, in some cases exceeding that of *dnaX*. It has been noted before that this pattern does not seem to be necessary for efficient -1 shifting^3^, and our results provide support for this. Efficient +1 sites are mainly characterized by a P-site allowing +1 rebinding of the cognate tRNA with at most one mismatch, and on the sites with two mismatches, one of them would be a G-U match, which is more tolerable than other non-Watson-Crick pairs^4,33,52^. In general, the P-site seems to be much more important for efficient frameshifting than the A-site, for both -1 and +1 shifting.

We note that our assay does not consider factors outside a short slippery site that influence frameshifting. These include mRNA secondary structures that interact with the ribosome^47,53,54^, longer codon repeats^31,55^, and, for prokaryotic systems, 5’ Shine-Dalgarno-like sequences that bind to the rRNA^56,57^. While complex interactions between the slippery site and the stimulators are likely, the relationship between codon patterns and FS probability that we observe, as well as previous work^3^, support our assumption that within a given context, frameshift is mainly thermodynamically controlled by the (re-)binding ability of the tRNAs.

Based on this assumption, we present a simple mechanistic model of frameshifting, SLIPPERRS, expanding on previous work modeling amino acid substitutions^33^. It is able to identify the most efficient frameshift sites with 73% accuracy, and requires no frameshift training data. It can be applied to any organism with an estimate of tRNA abundances and sufficient amino acid substitution data to fit nucleotide interaction parameters, for example by using deTELpy^40^. Several extensions to the model are conceivable, such as accounting for nucleotide modifications beyond lysidine, or allowing for stop codons. As it is, the model can be used to identify likely candidates for frameshift sites and to compare patterns of frameshift site occurrence across proteomes. SLIPPERS is the first general model for all types of FS sites: programmed or not, and backward (-1) and forward (+1), as previous models relied on *a priori* knowledge on frameshift sequence patterns and/or pre-defined mRNA secondary structure elements, e.g. pseudoknots (KnotInFrame^58^, FSFinder^59^), which may exclude novel or atypical frameshifting signals. PRFect, a recent machine learning model, considers many factors known to make FS more likely, but is only applicable to programmed frameshifts, and only considers a small, fixed set of slippery sites^38^.

Our dataset of FS sites enabled us to explore the extent of frameshifting more broadly, using *Gammaproteobacteria* as a model. It appears that frameshifting is a relevant factor in their genomes, based on two observations: First, efficient frameshift sites exist in a large number of genes, dozens per species on average. Second, frameshifts at the most efficient sites would produce smaller, more ordered proteins than expected. Larger proteins cost more energy to produce, while disordered proteins are more likely to be toxic (e.g., via aggregation), and prokaryotes in particular rarely make use of disordered protein regions^60,61^. It is possible that there has been selection to avoid producing wasteful, potentially deleterious proteins via frameshifting, for example via substitutions synonymous to the canonical frame, or by new cryptic (shifted-frame) stop codons. In addition, the difference between the products of high- and low-FS sites increases with the genomic GC content. The length of FS proteins is expected to increase with GC content, and the same holds for their disorder^62,63^. We observe the biggest difference in the length of frameshift products between high- vs. low-FS sites for species with high GC content, indicating that those species where the consequences of frameshifting would be most severe underwent the strongest selection to counteract them. In this context, it is noteworthy that the standard genetic code was previously shown to be close to optimal to reduce changes in proteins’ physicochemical properties upon frameshifting^64,65^.

Our results indicate that ribosomal frameshifting is a widespread phenomenon in bacteria and its impact is carefully tuned to mitigate deleterious effects, for example by selecting for cryptic stop codons. Our work enables further exploration of frameshifting in *Gammaproteobacteria* and beyond. We trust that both our dataset of experimentally tested FS sites and SLIPPERRS’ predictions will facilitate further study on the extent of ribosomal frameshifting and its impact on protein regulation and evolution.

## METHODS

### Frameshift model – SLIPPERRS

Our mechanistic model of frameshifting identifies 10-mer causing frameshifting by -2, -1, +1, and +2 based on three elements: The 10-nucleotide FS site itself, the abundances of tRNAs in the cell, and affinity parameters for codon-anticodon interaction. For the purpose of modeling, a frameshift happens if a tRNA is incorporated in the ribosomal A-site in a shifted frame. Both intuitively and based on current knowledge of the mechanism of frameshifting, this also requires the tRNA in the ribosome’s P-site to rebind to the new frame. A frameshift thus requires these two events to coincide. This may happen either during decoding or during translocation, and we model both these cases: A decoding shift happens if a tRNA arrives at the ribosome and binds to a shifted-frame codon in the A-site, and the P-site tRNA rebinds to the same frame. For a translocation shift, both P-site and A-site tRNA rebind together after binding of a tRNA in the A-site. Briefly, we first compute the energetic landscape of P-site rebinding and A-site binding separately, and then combine the two and apply the partition function to obtain binding probabilities. The binding probabilities are scaled by tRNA abundance. Finally, we calculate the probabilities of A-site and P-site rebinding and combine these with the scaled binding probabilities.

To calculate binding probabilities, we assume thermodynamic equilibrium, which means that the binding probability of a given tRNA to a codon is purely a function of the binding energy or affinity between the tRNA’s anticodon and the codon. In nature, this binding energy likely is influenced by the specific environment of the ribosome, forming many interactions with both the mRNA and the tRNA. Instead of trying to reflect this complex process, we calculate an *effective binding affinity* based on nucleotide interaction parameters and position-specific parameters fitted to observed amino acid substitutions in previous work^33^. The nucleotide interaction parameters are position-independent, but not symmetric, such that there is one parameter for every combination of codon nucleotide and anticodon nucleotide. The position parameters reflect the different degrees of ‘control’ the ribosome exerts on nucleotide binding at the three codon-anticodon positions. The binding affinity 𝛥𝐴 of codon *i* and anticodon *j* is then a linear combination of position-independent, non-symmetric parameters for the interaction of codon nucleotide *c* and anticodon nucleotide *ac* and a position-specific parameter *s* for each position *k*:

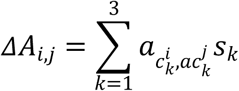

The first step in the model is to calculate this effective binding affinity for both P-site and A-site for all frames: -2, -1, 0, +1, +2. Even though -2/+1 or -1/+2 shifts are equivalent in that they lead to the same codon sequence, they are treated separately in the model, because they require the binding of different codons. For each site (P and A) there are thus five codons (or equivalently, frames) to consider.

For the case of P-site rebinding, we assume a correct tRNA, meaning cognate to the 0-frame codon, is already incorporated. In the case of several cognate tRNAs, we calculate their binding probabilities to the 0-frame codon by calculating the affinities and applying the partition function, as described below. We thus obtain an ‘expected’ tRNA, which is an average over all cognate tRNAs, represented as the set of their binding probabilities (summing to 1). We then calculate affinities of this expected tRNA to the shifted codons by calculating affinities of each possible tRNA, weighing them with their binding probability, and summing them. This results in one affinity value for each frame.

In the A-site, we allow the incorporation of any tRNA into any frame. We therefore calculate the effective binding affinities of all tRNAs to the 0-frame and all four shifted codons.

We then combine P-site and A-site affinities: We sum, for each frame, the affinities of each tRNA with the P-site affinity for that frame. We thus obtain a combined energy landscape, representing all possible events in which a tRNA binds to the A-site and the P-site tRNA rebinds accordingly.

We assume that it is not possible for the P-site tRNA and A-site tRNA to be bound to different frames. From this it follows that all binding events are mutually exclusive. Therefore, we calculate the binding probability of a tRNA to a codon by applying the partition function, using these affinities as pseudo-energies and summing over all tRNAs *m* and all codons *n* (or, equivalently, frames):

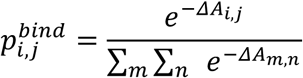

To obtain final incorporation probabilities, we need to account for differences in the abundance of tRNAs in the cell, as a more abundant tRNA will have more chances to bind to the ribosome and have its amino acid incorporated. In this work, we used tRNA abundance values for *E. coli* from ^66^. We simply scale the binding probabilities by the tRNA abundances *a* and normalize again:

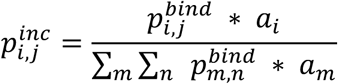

This calculation results in a matrix of incorporation probabilities for each tRNA in each frame, reflecting frameshifting during decoding.

To reflect the possibility of rebinding after tRNA incorporation, we make use of the same combined energy landscape described above. To this matrix of the shape (tRNAs x shift types), we apply the partition function row-wise, to each tRNA separately. This way we obtain conditional probabilities: The probability that P-site and A-site tRNA will rebind, given that the A-site has bound. Since we consider frameshifting during incorporation and frameshifting via rebinding independent events, we sum the incorporation and rebinding probabilities to acceptance probabilities, and normalize to obtain a proper probability space.

We obtain final scores by summing acceptance probabilities of all tRNAs carrying the same amino acid. The output is then a score between 0 and 1 for each amino acid in each frame, summing to 1. We treat the model output as a score indicating the probability to cause frameshifting. By applying a threshold, the output can be turned into a classification.

### Frameshift testing library

SLIPPERRS scores were calculated for all possible 10-mers (4^10^, approx. 10^6^ sequences), excluding those with a Stop codon in either the P-site or A-site. The list of predictions was filtered according to two criteria: No match to the canonical -1 pattern X_XXY_YYZ (where XXY is the P-site and YYZ the A-site; X, Y, and Z are each one nucleotide), and no stretch of more than four identical nucleotides. In addition, positive sequences needed to occur at least once in a canonical *E. coli* coding sequence (from NCBI RefSeq genome GCF_000005845.2) in the correct frame (such that P-site and A-site align with the 0-frame, 10-mer starts in +1- frame). From the remaining sequences, negative sequences were randomly selected from all 10-mers with a frameshift score of less than 0.001 for all frames. Positive sites were selected by randomly selecting among sequences with a score of at least 0.1 for the respective shift type. Across the library, sequences were required to have a minimum edit distance (Hamming distance, number of substitutions) of 3 to all other library sequences. For all four shift types considered here (-2, -1, +1, +2), both positive and negative predictions were selected (160 sequences in total, see supp. data). The 0-control sequence used as a reference for frameshift quantification was taken from Kelly et al. (2020)^45^, and additionally included in the -1 and +1 frames. Other control sequences included the release factor 2 site from *E. coli*, the *E. coli* dnaX site, and the SARS-CoV2 site, all of which were identical to the respective natural coding sequence, and a 10-fold thymine repeat (‘T-repeat’).

### Design of the frameshift testing construct

The protein sequence of the frameshift testing construct was designed by combining four basic components: The Q1 and Q2 peptides, the spacer regions before (N-terminal) and after (C-terminal) the FS site, and the mScarlet-I sequence.

The Q1 peptides were taken from Raghuraman et al. (2022)^42^, where they were used and referred to as reference peptides. Analogous to their approach, the Q2 peptides are modified versions of the Q1 peptides by exchanging one amino acid with similar physicochemical properties and by swapping two adjacent amino acids (see Table S1 for all peptides).

The mScarlet-I sequence is the original sequence designed in Bindels et al. (2017)^67^. Its coding sequence was codon-optimized for *E. coli* and not further modified.

The coding sequence for the Q-peptides and the spacer regions was codon optimized for *E. coli* using the GenSmart codon optimization tool (Genscript).

In order to avoid FS sites in the construct beyond the designed 10-mer, SLIPPERRS was applied to all 0-frame 10-mers in the coding sequence (one 10-mer per codon), and synonymous substitutions were made to remove all predicted frameshift sites. A single nucleotide insertion (for -1) or deletion (for +1) was made immediately after the 10-mer to shift the Q2 peptides and mScarlet-I into different frames relative to the start of the construct.

### Plasmids

The plasmid backbone used for all constructs was based on pEXP5-NT and contained the T7 promoter, the protein coding sequence, Broccoli tag, and Amp resistance gene (Figure S1). Plasmids containing the construct and the variable 10-mer sites were ordered from Genscript.

Lyophilized DNA was reconstituted in MilliQ water to a concentration of 100ng/µL and stored at -20°C. Selected sequences (e.g., controls) were amplified by transformation into chemically competent cells (XL1-blue, NEB), grown at 37°C in LB and the plasmids were purified using NucleoBond Xtra Midi Kit (Macherey-Nagel). After determining DNA concentration by absorption at 260nm (Nanodrop), purified stock solutions were stored at -20°C and diluted to 100ng/µL before use.

### *In vitro* protein synthesis reaction and frameshift measurement

Cell-free *in vitro* protein synthesis was carried out using the PURExpress transcription/translation system (New England Biolabs). The reaction mix was assembled as a master mix according to the manufacturer’s instructions, but at half volume: Per reaction, 5µL reagent A, 3.75 µL reagent B, 0.25 µL RNAse inhibitor murine (NEB), 1µL water (MilliQ) were used. DNA samples were at 100 ng/µL or approx. 40nM. Reactions were assembled in a 384 well-plate by adding 10µL reaction mix to each sample well, then adding 2.5µL sample for a final DNA concentration of 8nM. All samples were measured in triplicates. Wells surrounding the sample wells were filled with 10µL water to reduce evaporation and the plate was covered with adhesive film. The measurement was started within 15 minutes of assembling the reaction.

Fluorescence measurements were carried out using a Spark 20M microplate reader (Tecan). The temperature was maintained at 37°C. Excitation was at 564nm, detection at 609nm, both with a bandwidth of 20nm. Gain was fixed at 70, z-position was optimized using a fluorescence-positive sample. Fluorescence was measured at 10-minute intervals overnight.

To calculate relative fluorescence, only endpoint values were considered (at 16 hours). The mean fluorescence of negative control (reaction mix and water as sample) was subtracted from all samples. As positive control, we used a construct expressing mScarlet-I in the normal 0- frame control (’0-control’, mScarlet-I encoded in the 0-frame). Relative fluorescence was then calculated as the ratio of the mean of triplicates of a sample to the mean of triplicates of the positive control. The baseline originating from alternative initiation was subtracted from the relative fluorescence of all samples. Since frameshifting occurs in the positive control construct as well, we used the -1 and +1 constructs containing the same sequence to correct for that. Specifically, the relative fluorescence of 0-control and the -1 and +1 constructs containing the same 10-mer were summed to obtain the reference value. Frameshift probability was then calculated as the ratio of the relative fluorescence of a sample to the reference value.

### *in vivo* frameshift measurement

For transformation of bacteria, 20µL of chemically competent cells (*E. coli* strain BL23 DE(pLys), Agilent) were thawed on ice. 0.5µL of DNA (100ng/µL) were added and the mix was incubated on ice for 30min. A heat shock was applied using a water bath at 42°C for 45s. Cells were put back on ice for 2min, then 500µL of prewarmed SOC medium was added. Cells were incubated at 37°C for 1h in a thermoshaker. 200µL of the mix were plated on LB agar containing 100µg/mL ampicillin. Plates were incubated at 37°C overnight.

For each sequence (10-mer), three independent colonies were used to inoculate three cultures (biological replicates) in 5mL of LB medium (100µg/mL Amp, 30µg/mL chloramphenicol). Cells were grown in deep-well plates overnight at 37°C, shaking at 180rpm.

The overnight precultures were diluted 1:50 with antibiotic-free LB medium. Cultures were then grown at 37°C, 180rpm shaking. After 3h, protein production was induced by adding Isopropyl β-d-1-thiogalactopyranoside (IPTG) to a concentration of 200µM.

Aliquots for measuring fluorescence were taken after 16h post-induction. 100µL of culture were measured in a 96-well plate (Greiner) using a Tecan Spark 20M plate reader. mScarlet-I fluorescence was measured by excitation at 564nm and detection at 609nm, bandwidth 20nm, fixed gain 100. Frameshift probabilities were calculated analogous to in-vitro experiments: The mean fluorescence of negative control was subtracted from all samples. Relative fluorescence was then calculated as the ratio of the mean of triplicates of a sample to the mean of triplicates of the positive control. As a negative control we used bacteria transformed with an empty vector (pUC18), and as a positive control we used bacteria transformed with the 0-control plasmid.

### Frameshift quantification by mass spectrometry

After *in vitro* protein synthesis reaction, triplicates were pooled and samples stored at -70°C. For in-solution digestion, RapiGest SF (Waters) in 50mM NH4HCO3 was added to samples to a final concentration of 0.15%. Protein concentration was determined by BCA assay (Pierce BCA Protein Assay Kit, Thermo Fisher Scientific). Disulfide bonds were reduced by adding dithiothreitol to a final concentration of 5mM and incubation at 37°C for 30min. Proteins were alkylated by adding iodoacetamide to a final concentration of 15mM and incubation at room temperature for 30min in the dark. Proteases (trypsin/LysC mix, mass spectrometry grade, Promega) were added at a protease:protein ratio of 1:20 and incubated at 37°C overnight. After digestion, RapiGest was cleaved by adding trifluoroacetic acid to a final concentration of 0.5% and incubation at 37°C for 30min. Cleavage products were removed by centrifuging at 16,000g for 10min and the supernatant was collected. The supernatant was dried in a vacuum centrifuge and stored at -20°C. For injection, dried peptides were reconstituted in 5% formic acid for a final concentration of ca. 120ng/µL. 5µL (600ng peptides) were injected for analysis.

All LC-MS/MS analysis was performed on a nanoUHPLC Vanquish system interfaced on-line to an Orbitrap HF hybrid mass spectrometer (both Thermo Fisher Scientific). The nano-LC system was equipped with an Acclaim PepMap 100 75µm x 2cm trapping column and a 50cm µPAC analytical column (both Thermo Fisher Scientific).

Quantitative peptides (Q-peptides and mScarlet-I) were analyzed with data-independent acquisition (DIA). Peptides were separated using a two-step gradient: 20min 0-17.5% B, 20min 17.5%-35% (A: 0.1% aqueous formic acid, B: 0.1% formic acid in acetonitrile). MS settings for DIA were: MS1; 60,000 resolution, AGC 3e6, max. injection time 60ms, mass range 380- 1,300. MS2; 30,000 resolution, AGC 2e5, automatic max. injection time, mass range 380- 1,200, window width 25m/z, normalized collision energy 28. Windows were placed with a 0.5m/z overlap on both sides.

DIA raw files were first converted to mzML with ProteoWizard ^68^. mzML files were analyzed with DIA-NN 1.8^69^ in library-free mode: First, a spectral library was generated in-silico using a fasta file containing the construct and predicted frameshift sequences, the canonical *E. coli* K-12 MG1655 proteome (NCBI) and common contaminants ^70^. The library was generated with the following settings: Cut at K/R, up to two missed cleavages, peptide length 7-50, precursor m/z 300-1,200, precursor charge 1-4, C(carbamidomethylation) as fixed modification, M(oxidation) as variable modification. Peptides were then identified and quantified with the following settings: Precursor and protein FDR 1%, double search enabled, isoleucine/leucine treated equivalent, smart library profiling, no interference removal. Frameshift probabilities were determined using the MS1-based precursor quantities of the Q1 and Q2 peptides as given in the DIA-NN main output (‘Ms1.Translated’). Precursor intensities of the same peptide were summed. The Q-peptide pair DAFIGTFLYEYSR/DAFIGTFYLDYSR was not reliably detected and was excluded from quantification. The abundance ratio of each peptide pair was calculated as the ratio of intensities of the respective Q2 and Q1 peptide. The frameshift probability was then calculated as the median of Q-peptide abundance ratios.

mScarlet-I was quantified using the peptides LYPEDGVLK and LDITSHNEDYTVVEQYER. MS1 intensities for these peptides were used as described above. The precursor intensities of these peptides were summed to obtain an mScarlet-I intensity value that was then comparable across samples.

### Identification of frameshift peptides

Identification of candidate frameshift peptides (peptides covering the frameshift site) was done using the same samples as the quantification. Sample preparation and mass spectrometry instrumentation were identical to the quantitative measurements described above, Samples were analyzed with targeted acquisition (PRM) using an inclusion list. The inclusion list was specific to the analyzed sample and contained predicted peptides resulting from frameshifts of different types and at slightly varying positions (10-mer’s A-site codon and surrounding codons). Peptides were monitored across the full gradient, without scheduling.

Peptides were separated with a 60-minute gradient 5-35% B at a flow rate of 0.5μL/min. A full mass spectrum was acquired over the range 350-1,700m/z with 60,000 resolution, AGC 3e6, max. injection time 60ms. MS2 scans for each peptide in the inclusion list were performed with 15,000 resolution, AGC 1e5, max. injection time 50ms, normalized collision energy 25, isolation window width 1.6m/z, and fixed first mass 140m/z.

Raw files from PRM measurements were converted to mzML using ProteoWizard. mzML files were analyzed using Skyline daily v22.2^71^: A fasta containing predicted frameshift sequences specific to the analyzed sample was loaded into Skyline. The list of precursors generated by Skyline was filtered for redundancy. For each relevant precursor (frameshift peptide), the correct peak was manually identified based on the presence and agreement of fragment ions and agreement between replicates and peptides of a Q-peptide pair. Peptide reports containing identification and quantification (MS1-based) were exported from Skyline for further analysis.

### Estimation of alternative initiation using absolute quantification of frameshift testing construct

A gel sample containing BSA was prepared by diluting NIST-certified BSA standard solution to 3pmol/µL. This was 1:2 with sample buffer (2x Laemmli buffer with 0.1M dithiothreitol).

6.7µL of the solution (corresponding to 10pmol/lane) were loaded in four replicates onto a NuPage 4-12% Bis-Tris gel. The gel was run in MOPS buffer at 200V for 40min. The gel was briefly rinsed in water, stained using ReadyBlue and destained in water. The gel regions containing BSA were cut and stored at -20°C.

To prepare samples after *in vitro* protein synthesis reaction, triplicates were pooled and mixed 1:1 with sample buffer. 6µL sample (corresponding to 3µL cell-free expression reaction mix) were loaded on to a 4-12% NuPage Bis-Tris gel. 1.5µL of purified mScarlet-I (100µg/mL) were mixed 1:1 with sample buffer and loaded on a separate lane as a weight marker. The gel was run in MOPS buffer at 200V for 40min, stained using ReadyBlue and destained in water. From the gel, a region corresponding to a molecular weight of ca. 20-45kDa was cut (Figure S5). Crucially, this region contains the full-length frameshift product (42kDa), as well as any products of alternative initiation that resulted in the production of full-length mScarlet-I (26- 37kDa), but not products of canonical translation terminated shortly after the FS site, <10kDa). The selected gel pieces were prepared for MS analysis together with one aliquot (one gel piece) of BSA: The gel pieces were cut into 1-2mm cubes and washed with 0.1M ammonium bicarbonate buffer (Ambic). The gel pieces were then shrunk by adding acetonitrile, shaking intensely for 1-2min, shortly centrifuging and removing the supernatant. Disulfide bonds were reduced by reconstituting the shrunken pieces in Ambic buffer containing 5mM DTT and incubation for 30min at 37°C. The gel pieces were then shrunk again. For alkylation, the pieces were reconstituted in Ambic buffer containing 15mM iodoacetamide and incubated at 30min at room temperature in the dark. The pieces were then shrunk, washed once with Ambic, and shrunk again. For enzymatic digestion, trypsin (MS grade, Promega) was dissolved to a concentration of 13.3mg/L in 40mM Ambic buffer containing 10% acetonitrile and added to the gel pieces to reconstitute. Samples were incubated for ca. 1h on ice and then at 37°C overnight.

To extract peptides after digestion, gel pieces were shrunk again and the supernatant collected. The pieces were reconstituted in 5% formic acid (FA) in water, shrunk, and the supernatant collected. The samples were shrunk again and the supernatant collected. All supernatants were pooled and dried down in a vacuum centrifuge.

Before MS analysis, BSA was reconstituted in 100µL 5% FA. Samples were reconstituted in 35µL 5% FA and 1µL of the BSA digest was added (for a final concentration of 2.8fmol/µL BSA). 5µL were injected into the LC-MS system for analysis. Peptides were separated using a two-step gradient: 20min 0-17.5% B, 20min 17.5%-35%. Peptides were analyzed using data-dependent acquisition with the following settings: MS1: 60,000 resolution, AGC target 3e6, max. injection time 60ms, scan range 250-1,600Th. MS2: Top 20 precursors analyzed, 15,000 resolution, AGC target 1e5, max. injection time 1e5, isolation window 1.6Th, normalized collision energy 25.

Raw files were converted to mzML using ProteoWizard. mzML files were analyzed using Skyline daily v22.2^71^: A fasta containing sequences of the frameshift testing construct, the *E. coli* genome, and contaminants was loaded into Skyline. The list of precursors generated by Skyline was filtered for redundancy. For each relevant precursor (see Table S1), the correct peak was manually identified based on the presence and agreement of fragment ions and agreement between replicates and peptides of a Q-peptide pair. Peptide reports containing identification and quantification (MS1-based) were exported from Skyline for further analysis. Absolute quantification of Q-peptides was based on the fact that the Q-peptides are slightly mutated versions of BSA peptides, allowing direct comparison of their MS signals. The peak areas (intensities) of all Q-peptides and the corresponding BSA peptides were collected. The amounts of Q1 and Q2 were then calculated by calculating the ratio of their intensity to the intensity of the corresponding BSA peptide.

To determine the baseline caused by alternative initiation, we make use of the fact that our samples only include full-length frameshift product (containing Q1, Q2, and mSc) and any products of alternative initiation (containing mSc, but not Q1). Any fluorescence measured in the absence of Q1 is then due to alternative initiation. We plotted the amounts of Q1 against relative fluorescence for all included samples and calculated a linear regression. The intercept of the regression line with the x-axis (Q1 = 0) is the baseline caused by alternative initiation.

### Codon analysis and sequence properties

The classification of P-site and A-site codons as ‘rare’ was based on relative synonymous codon usage (RSCU). To calculate RSCU for *E. coli*, integrated protein abundances were retrieved from paxDB^72^ (‘whole organism - Integrated’ dataset, retrieved 19/04/2023) and the top 5% proteins by abundance were selected. From the coding sequences of these proteins, the occurrences of each codon were counted. The RSCU of a codon X is then the ratio of the frequency of X and the mean frequency of all codons synonymous to X (including X). All codons with an RSCU < 1 were classified as ‘rare’.

A codon was classified as having a ‘perfect’ tRNA if the tRNA pool of *E. coli* contains a tRNA with an anticodon able to form three Watson-Crick pairings with the codon.

To define whether a sequence has a ‘slippery’ A- or P-site, we determined whether the tRNA cognate to the 0-frame codon would also bind in the shifted frame. The criteria for this were based on^4^, and the classification was done separately for -1 and +1 shifting for each sequence: For a given shift type, 10-mer, and site (P or A), we looked at the pairing between anticodon and the shifted codon. The site was considered slippery in the following case: Among the first two codon positions, at least one had to be a Watson-Crick pairing, and the other either Watson-Crick or a GU pairing, which is also considered slightly favorable^4,33^. We allowed a mismatch in the third (wobble) position. The assignment of which codon is decoded by which tRNA/anticodon was retrieved from Avcilar-Kucukgoze et al. (2016)^73^. If several anticodons are known to decode a given codon, we only required one of them to be able to rebind to consider the site slippery.

The calculation of relative risk was based on a 2x2 contingency table. The table would contain counts of: pp = 10-mers with the property and predicted positive, p = total 10-mers with the property, ap = 10-mers without the property and predicted positive, a = total 10-mers without the property. The relative risk is then (pp / p) / (ap / a). The risk ratio and 95% confidence intervals were calculated using the respective functions in SciPy 1.12.0 ^74^.

For the calculation of relative risk, 10-mers were included twice, once for each shift type. A single sequence could thus count as a positive and a negative, as two positives, or two negatives, if it is predicted to shift by +1, but not -1, both, or neither, respectively.

### Mapping of frameshift sites and analysis of hypothetical frameshift proteins

To identify potential frameshift sites, the occurrences of all tested 10-mers in the coding sequences of the 200 *Gammaproteobacteria* species were enumerated. 10-mers had to be in the correct frame, such that P-site and A-site codon are in 0-frame (start of the 10-mer in +1 frame), and fully within the coding sequence. To generate hypothetical frameshift proteins, we first retrieved, for each gene, an ‘extended CDS’ containing the first 1,500 nucleotides 3’ of the gene’s Stop codon (based on the annotated genomic position), accounting for both the strand on which the gene was located and the circularity of the genome. This was done for all genes regardless of the occurrence of other genes, pseudogenes or other genomic elements in this region. Cases where the frameshifted sequence would ‘fuse’ with a different gene were annotated and excluded from all analyses concerning the shifted protein sequences. Frameshifted protein sequences were obtained by simply translating the extended coding sequence, with the respective shift at the first position of the 10-mer’s A-site codon.

10-mers with an FS probability of at least 10.4% were classified as ‘high-FS’, all others as ‘low-FS’.

Protein disorder was predicted using IUPred2A ‘long’^75^. Positions with an IUPred score >= 0.5 were annotated as disordered.

Pfam domains were annotated using InterProScan 5.73-104.0^76,77^ locally with standard options. Protein 3D structures were predicted with the Alphafold webserver (Alphafold3^78^).

For categorization of frameshift consequences, for each frameshift product, Pfam domains were annotated in the frameshifted protein and the canonical protein. A domain identified in the canonical sequence was considered ‘preserved’ if the same Pfam domain was identified in the frameshift sequence with at least 80% of the original domain’s length and starting within 10 residues of the original start, otherwise the domain was considered ‘lost’. The frameshift’s consequence was then categorized as follows: ‘Destroyed’, none of the domains in the canonical sequence were preserved; ‘Truncated’, at least one of the canonical domains was preserved; ‘Preserved’, all canonical domains were preserved. If the frameshift sequence contained a domain with a start site after the frameshift region, the protein was classified as ‘Frameshift domain’. Each protein would thus be classified as either ‘Destroyed’, ‘Truncated’, or ‘Preserved’, and might additionally be classified to have a ‘Frameshift domain’. ‘Fused’ cases (where the frameshift leads to a fusion with a second gene) were considered a separate category and excluded from the other categories.

### Statistical and data analyses

All scripting, data processing and statistical analysis was done using custom code written in Python 3.9 or 3.10.^79^ and Jupyter Notebooks, with the help of Numpy 1.23^80^ and pandas 2.2^81^. Statistical tests and model performance evaluations were done using the corresponding functions in Scipy 1.12^74^ and scikit-learn 1.2^82^. Sequence data was processed with the help of Biopython 1.79^83^. Protein structures were visualized using Pymol^84^.

For all boxplots, boxes indicate the 25-75 percentile, whiskers extend to 1.5x the interquartile range, and outliers beyond this range are displayed as points.

## AUTHOR CONTRIBUTIONS

A.T-P., Andrej S. and J.P. conceived the project and designed experiments. J.P. and C.L. developed the computational model. J.P. performed experiments with the assistance of D.R. and Anna S. D.R. and J.P. performed *in vivo* experiments. M.L.R.R. provided reagents and designed experiments. J.P. and A.T-P. performed formal data analysis. J.P. prepared all the figures and wrote the initial draft of the manuscript. J.P. and A.T-P. prepared the final manuscript with input from all authors. A.T-P. and Andrej S. acquired funding.

## ACKNOWLEDGEMENTS

We would like to thank members of the Shevchenko lab, especially Tobias Jumel, Ignacy Rzagalinski, Viditha Rao, and Andrea Schuhmann for the help and advice with mass spectrometry. We thank members of the Toth-Petroczy lab for helpful discussions, and especially Anna Hadarovich for her support with Alphafold2, Maxim Scheremetjew for the assistance with code documentation and publication, and Federica Luppino for her advice on statistical analysis. We would like to thank the Computer Service of the MPI-CBG for their support, especially to Oscar Gonzales for supporting our HPC. We are grateful to Rico Barsacchi and Martin Stöter of the Technology Development Studio of the MPI-CBG for support with the *in vitro* assay. We thank Eric Geertsma and Aliona Bogdanova from the Protein Biochemistry facility of the MPI-CBG for providing the expression vector and for their support with protein design and expression.

Figures 2A and 6A were created with the help of biorender.com.

## FUNDING

This work was funded by the Max Planck Gesellschaft (MPG).

## CONFLICT OF INTEREST

The authors declare no conflicts of interest.

## Notes

### Competing Interest Statement

The authors have declared no competing interest.

## References

1. Romero Romero, M. L., Landerer, C., Poehls, J. & Toth-Petroczy, A. Phenotypic mutations contribute to protein diversity and shape protein evolution. Protein Sci. 31, (2022).

2. Sharma, V. et al. Analysis of tetra- and hepta-nucleotides motifs promoting -1 ribosomal frameshifting in Escherichia coli. Nucleic Acids Res. 42, 7210–7225 (2014).

3. Bock, L. V. et al. Thermodynamic control of −1 programmed ribosomal frameshifting. Nat. Commun. 10, 1–11 (2019).

4. Springstein, B. L. et al. Systematic analysis of nonprogrammed frameshift suppression in E. coli via translational tiling proteomics. Proc. Natl. Acad. Sci. U. S. A. 121, e2317453121 (2024).

5. Atkins, J. F., Loughran, G., Bhatt, P. R., Firth, A. E. & Baranov, P. V. Ribosomal frameshifting and transcriptional slippage: From genetic steganography and cryptography to adventitious use. Nucleic Acids Res. 44, 7007–7078 (2016).

6. Farabaugh, P. J. Programmed Frameshifting in Budding Yeast. in Recoding: Expansion of Decoding Rules Enriches Gene Expression (eds. Atkins, J. F. & Gesteland, R. F.) 221–247 (Springer New York, New York, NY, 2010).

7. Caliskan, N., Katunin, V. I., Belardinelli, R., Peske, F. & Rodnina, M. V. Programmed -1 frameshifting by kinetic partitioning during impeded translocation. Cell 157, 1619–1631 (2014).

8. Brierley, I., Gilbert, R. J. C. & Pennell, S. Pseudoknot-dependent programmed —1 ribosomal frameshifting: Structures, mechanisms and models. in *Recoding: Expansion of Decoding Rules Enriches Gene Expression* 149–174 (Springer New York, New York, NY, 2010).

9. Demo, G. et al. Structural basis for +1 ribosomal frameshifting during EF-G-catalyzed translocation. Nat. Commun. 12, 4644 (2021).

10. Korniy, N. et al. Modulation of HIV-1 Gag/Gag-Pol frameshifting by tRNA abundance. Nucleic Acids Res. 47, 5210–5222 (2019).

11. Caliskan, N. et al. Conditional Switch between Frameshifting Regimes upon Translation of dnaX mRNA. Mol. Cell 66, 558–567.e4 (2017).

12. Kramer, E. B. & Farabaugh, P. J. The frequency of translational misreading errors in E. coli is largely determined by tRNA competition. RNA 13, 87–96 (2007).

13. Dulude, D., Baril, M. & Brakier-Gingras, L. Characterization of the frameshift stimulatory signal controlling a programmed –1 ribosomal frameshift in the human immunodeficiency virus type 1. Nucleic Acids Res. 30, 5094–5102 (2002).

14. Plant, E. P., Rakauskaite, R., Taylor, D. R. & Dinman, J. D. Achieving a golden mean: mechanisms by which coronaviruses ensure synthesis of the correct stoichiometric ratios of viral proteins. J. Virol. 84, 4330–4340 (2010).

15. Lobanov, A. V. et al. Position-dependent termination and widespread obligatory frameshifting in Euplotes translation. Nat. Struct. Mol. Biol. 24, 61–68 (2017).

16. Ivanov, I. P. & Matsufuji, S. Autoregulatory Frameshifting in Antizyme Gene Expression Governs Polyamine Levels from Yeast to Mammals. in Recoding: Expansion of Decoding Rules Enriches Gene Expression (eds. Atkins, J. F. & Gesteland, R. F.) 281–300 (Springer New York, New York, NY, 2010).

17. Russell, R. D. & Beckenbach, A. T. Recoding of translation in turtle mitochondrial genomes: programmed frameshift mutations and evidence of a modified genetic code. J. Mol. Evol. 67, 682–695 (2008).

18. Yanagida, H. et al. The Evolutionary Potential of Phenotypic Mutations. PLoS Genet. 11, e1005445 (2015).

19. Wills, N. M., Moore, B., Hammer, A., Gesteland, R. F. & Atkins, J. F. A functional -1 ribosomal frameshift signal in the human paraneoplastic Ma3 gene. J. Biol. Chem. 281, 7082–7088 (2006).

20. Asakura, T. et al. Isolation and characterization of a novel actin filament-binding protein from Saccharomyces cerevisiae. Oncogene 16, 121–130 (1998).

21. Manktelow, E., Shigemoto, K. & Brierley, I. Characterization of the frameshift signal of Edr, a mammalian example of programmed -1 ribosomal frameshifting. Nucleic Acids Res. 33, 1553–1563 (2005).

22. Tsuchihashi, Z. & Kornberg, A. Translational frameshifting generates the gamma subunit of DNA polymerase III holoenzyme. Proc. Natl. Acad. Sci. U. S. A. (1990).

23. Meydan, S. et al. Programmed Ribosomal Frameshifting Generates a Copper Transporter and a Copper Chaperone from the Same Gene. Mol. Cell 65, 207–219 (2017).

24. Craigen, W. J. & Caskey, C. T. Expression of peptide chain release factor 2 requires high-efficiency frameshift. Nature 322, 273–275 (1986).

25. Baranov, P. V., Gesteland, R. F. & Atkins, J. F. Release factor 2 frameshifting sites in different bacteria. EMBO Rep. 3, 373–377 (2002).

26. Jacks, T. et al. Characterization of ribosomal frameshifting in HIV-1 gag-pol expression. Nature 331, 280– 283 (1988).

27. Blinkowa, A. L. & Walker, J. R. Programmed ribosomal frameshifting generates the Escherichia coli DNA polymerase III gamma subunit from within the tau subunit reading frame. Nucleic Acids Res. 18, 1725– 1729 (1990).

28. Atkins, J. F., Elseviers, D. & Gorini, L. Low Activity of β-Galactosidase in Frameshift Mutants of Escherichia coli. Proc. Natl. Acad. Sci. U. S. A. 69, 1192–1195 (1972).

29. Kurland, C. G. Translational accuracy and the fitness of bacteria. Annu. Rev. Genet. 26, 29–50 (1992).

30. Marczinke, B., Hagervall, T. & Brierley, I. The Q-base of asparaginyl-tRNA is dispensable for efficient -1 ribosomal frameshifting in eukaryotes. J. Mol. Biol. 295, 179–191 (2000).

31. Ren, G. et al. Ribosomal frameshifting at normal codon repeats recodes functional chimeric proteins in human. Nucleic Acids Res. 52, 2463–2479 (2024).

32. Drummond, D. A. & Wilke, C. O. The evolutionary consequences of erroneous protein synthesis. Nat. Rev. Genet. 10, 715–724 (2009).

33. Landerer, C., Pöhls, J. & Toth-Petroczy, A. Fitness effects of phenotypic mutations at proteome-scale reveal optimality of translation machinery. Mol. Biol. Evol. 41, (2024).

34. Antonov, I., Coakley, A., Atkins, J. F., Baranov, P. V. & Borodovsky, M. Identification of the nature of reading frame transitions observed in prokaryotic genomes. Nucleic Acids Res. 41, 6514–6530 (2013).

35. Firth, A. E. Mapping overlapping functional elements embedded within the protein-coding regions of RNA viruses. Nucleic Acids Res. 42, 12425–12439 (2014).

36. Richardson, M. O. & Eddy, S. R. ORFeus: A Computational Method to Detect Programmed Ribosomal Frameshifts and Other Non-Canonical Translation Events. bioRxiv 2023.04.24.538127 (2023) doi:10.1101/2023.04.24.538127.

37. Michel, A. M. et al. Observation of dually decoded regions of the human genome using ribosome profiling data. Genome Res. 22, 2219–2229 (2012).

38. McNair, K., Salamon, P., Edwards, R. A. & Segall, A. M. PRFect: a tool to predict programmed ribosomal frameshifts in prokaryotic and viral genomes. BMC Bioinformatics 25, 82 (2024).

39. Mikl, M., Pilpel, Y. & Segal, E. High-throughput interrogation of programmed ribosomal frameshifting in human cells. Nat. Commun. 11, (2020).

40. Landerer, C., Scheremetjew, M., Moon, H., Hersemann, L. & Toth-Petroczy, A. deTELpy: Python package for high-throughput detection of amino acid substitutions in mass spectrometry datasets. Bioinformatics 40, (2024).

41. Rzagalinski, I. et al. FastCAT Accelerates Absolute Quantification of Proteins Using Multiple Short Nonpurified Chimeric Standards. J. Proteome Res. 21, 1408–1417 (2022).

42. Raghuraman, B. K. et al. Median-Based Absolute Quantification of Proteins Using Fully Unlabeled Generic Internal Standard (FUGIS). J. Proteome Res. 21, 132–141 (2022).

43. Fages-Lartaud, M., Tietze, L., Elie, F., Lale, R. & Hohmann-Marriott, M. F. mCherry contains a fluorescent protein isoform that interferes with its reporter function. Front. Bioeng. Biotechnol. 10, 892138 (2022).

44. Xu, J., Hendrix, R. W. & Duda, R. L. Conserved translational frameshift in dsDNA bacteriophage tail assembly genes. Mol. Cell 16, 11–21 (2004).

45. Kelly, J. A. et al. Structural and functional conservation of the programmed -1 ribosomal frameshift signal of SARS coronavirus 2 (SARS-CoV-2). J. Biol. Chem. 295, 10741–10748 (2020).

46. Yan, S., Wen, J.-D., Bustamante, C. & Tinoco, I., Jr. Ribosome excursions during mRNA translocation mediate broad branching of frameshift pathways. Cell 160, 870–881 (2015).

47. Lin, Z., Gilbert, R. J. C. & Brierley, I. Spacer-length dependence of programmed-1 or-2 ribosomal frameshifting on a U6A heptamer supports a role for messenger RNA (mRNA) tension in frameshifting. Nucleic Acids Res. 40, 8674–8689 (2012).

48. Curran, J. F. & Yarus, M. Use of tRNA suppressors to probe regulation of Escherichia coli release factor 2. J. Mol. Biol. 203, 75–83 (1988).

49. Zhang, D. et al. EF4 disengages the peptidyl-tRNA CCA end and facilitates back-translocation on the 70S ribosome. Nat. Struct. Mol. Biol. 23, 125–131 (2016).

50. Rajkovic, A. & Ibba, M. Elongation factor P and the control of translation elongation. Annu. Rev. Microbiol. 71, 117–131 (2017).

51. Saito, K. et al. Ribosome collisions induce mRNA cleavage and ribosome rescue in bacteria. Nature 603, 503–508 (2022).

52. Grosjean, H. J., de Henau, S. & Crothers, D. M. On the physical basis for ambiguity in genetic coding interactions. Proc. Natl. Acad. Sci. U. S. A. 75, 610–614 (1978).

53. Bhatt, P. R. et al. Structural basis of ribosomal frameshifting during translation of the SARS-CoV-2 RNA genome. Science 372, 1306–1313 (2021).

54. Mouzakis, K. D., Lang, A. L., Vander Meulen, K. A., Easterday, P. D. & Butcher, S. E. HIV-1 frameshift efficiency is primarily determined by the stability of base pairs positioned at the mRNA entrance channel of the ribosome. Nucleic Acids Res. 41, 1901–1913 (2013).

55. Girstmair, H. et al. Depletion of Cognate Charged Transfer RNA Causes Translational Frameshifting within the Expanded CAG Stretch in Huntingtin. Cell Rep. 3, 148–159 (2013).

56. Kim, H.-K. & Tinoco, I., Jr. EF-G catalyzed translocation dynamics in the presence of ribosomal frameshifting stimulatory signals. Nucleic Acids Res. 45, 2865–2874 (2017).

57. Chen, J. et al. Dynamic pathways of -1 translational frameshifting. Nature 512, 328–332 (2014).

58. Theis, C., Reeder, J. & Giegerich, R. KnotInFrame: prediction of -1 ribosomal frameshift events. Nucleic Acids Res. 36, 6013–6020 (2008).

59. Moon, S., Byun, Y., Kim, H.-J., Jeong, S. & Han, K. Predicting genes expressed via −1 and +1 frameshifts. Nucleic Acids Res. 32, 4884–4892 (2004).

60. Basile, W., Salvatore, M., Bassot, C. & Elofsson, A. Why do eukaryotic proteins contain more intrinsically disordered regions? PLoS Comput. Biol. 15, e1007186 (2019).

61. Chow, C. F. W. & Toth-Petroczy, A. The evolution and exploration of intrinsically disordered and phase-separated protein states. in The Three Functional States of Proteins 353–379 (Elsevier, 2025).

62. Pavlović-Lažetić, G. M. et al. Bioinformatics analysis of disordered proteins in prokaryotes. BMC Bioinformatics 12, 66 (2011).

63. Basile, W., Sachenkova, O., Light, S. & Elofsson, A. High GC content causes orphan proteins to be intrinsically disordered. PLoS Comput. Biol. 13, e1005375 (2017).

64. Xu, H. & Zhang, J. On the Origin of Frameshift-Robustness of the Standard Genetic Code. Mol. Biol. Evol. 38, 4301–4309 (2021).

65. Bartonek, L., Braun, D. & Zagrovic, B. Frameshifting preserves key physicochemical properties of proteins. Proc. Natl. Acad. Sci. U. S. A. 117, 5907–5912 (2020).

66. Larson, M. H. et al. A pause sequence enriched at translation start sites drives transcription dynamics in vivo. Science 344, 1042–1047 (2014).

67. Bindels, D. S. et al. mScarlet: a bright monomeric red fluorescent protein for cellular imaging. Nat. Methods 14, 53–56 (2017).

68. Adusumilli, R. & Mallick, P. Data Conversion with ProteoWizard msConvert. Methods Mol. Biol. 1550, 339–368 (2017).

69. Demichev, V., Messner, C. B., Vernardis, S. I., Lilley, K. S. & Ralser, M. DIA-NN: neural networks and interference correction enable deep proteome coverage in high throughput. Nat. Methods 17, 41–44 (2020).

70. Frankenfield, A. M., Ni, J., Ahmed, M. & Hao, L. Protein Contaminants Matter: Building Universal Protein Contaminant Libraries for DDA and DIA Proteomics. J. Proteome Res. 21, 2104–2113 (2022).

71. Egertson, J. D., MacLean, B., Johnson, R., Xuan, Y. & MacCoss, M. J. Multiplexed peptide analysis using data-independent acquisition and Skyline. Nat. Protoc. 10, 887–903 (2015).

72. Huang, Q., Szklarczyk, D., Wang, M., Simonovic, M. & von Mering, C. PaxDb 5.0: Curated protein quantification data suggests adaptive proteome changes in yeasts. Mol. Cell. Proteomics 22, 100640 (2023).

73. Avcilar-Kucukgoze, I. et al. Discharging tRNAs: a tug of war between translation and detoxification in Escherichia coli. Nucleic Acids Res. 44, 8324–8334 (2016).

74. Virtanen, P. et al. SciPy 1.0: fundamental algorithms for scientific computing in Python. Nat. Methods 17, 261–272 (2020).

75. Mészáros, B., Erdős, G. & Dosztányi, Z. IUPred2A: context-dependent prediction of protein disorder as a function of redox state and protein binding. Nucleic Acids Res. 46, W329–W337 (2018).

76. Jones, P. et al. InterProScan 5: genome-scale protein function classification. Bioinformatics 30, 1236–1240 (2014).

77. Paysan-Lafosse, T. et al. InterPro in 2022. Nucleic Acids Res. 51, D418–D427 (2023).

78. Abramson, J. et al. Accurate structure prediction of biomolecular interactions with AlphaFold 3. Nature 630, 493–500 (2024).

79. VanRossum, G. & Drake, F. L. The python language reference. https://scicomp.ethz.ch/public/manual/Python/3.9./reference.pdf.

80. Harris, C. R. et al. Array programming with NumPy. Nature 585, 357–362 (2020).

81. Pandas Team. Pandas-Dev/Pandas: Pandas. (Zenodo, 2024). doi:10.5281/zenodo.10697587.

82. Pedregosa, F. et al. Scikit-learn: Machine Learning in Python. Journal of Machine Learning Research 12, 2825–2830 (2011).

83. Cock, P. J. A. et al. Biopython: freely available Python tools for computational molecular biology and bioinformatics. Bioinformatics 25, 1422–1423 (2009).

84. Schrödinger, LLC. The PyMOL Molecular Graphics System, Version 1.8. Preprint at (2015).

85. Brademan, D. R., Riley, N. M., Kwiecien, N. W. & Coon, J. J. Interactive peptide spectral annotator: A versatile web-based tool for proteomic applications. Mol. Cell. Proteomics 18, S193–S201 (2019).

