## Supplementary Information for "Ribosomal frameshifting is a carefully tuned modifier of proteins"

**Figures S1-S11**

**Tables S1-S3**

**Supplemental References**

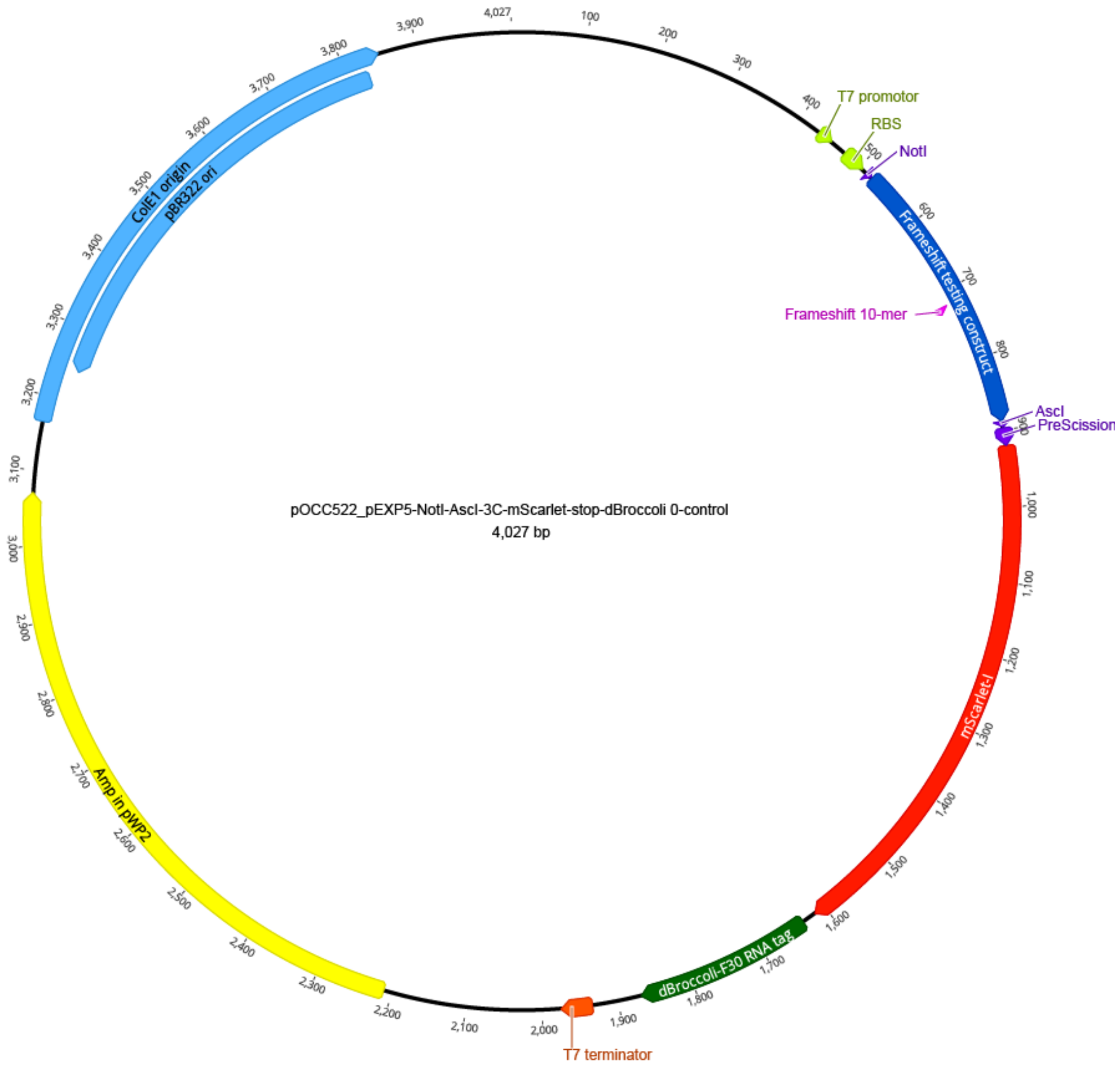


**Figure S1. Map of the plasmid used for frameshift quantification.**

The plasmid is based on pEXP5-NT. It contains an Ampicillin resistance gene, a Broccoli mRNA tag, the T7 promoter and terminator, and the coding sequence of mScarlet-I. The coding sequence of the rest of the frameshift testing construct (Q-peptides, frameshift site, spacers) was inserted 5‘ of mScarlet-I.

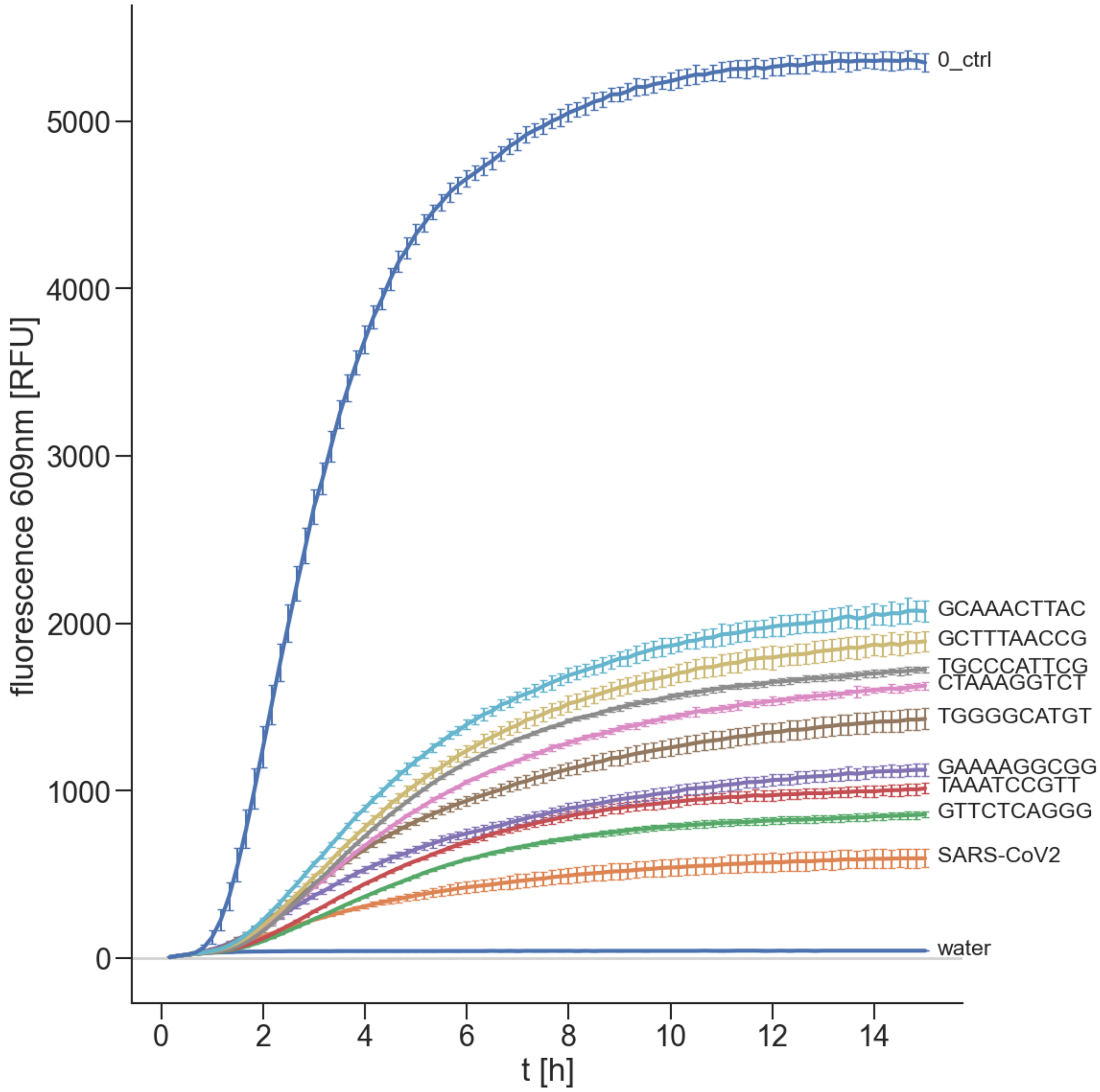


**Figure S2. Example raw fluorescence data for frameshift quantification.**
The cell-free reaction is carried out and monitored over up to 16h. mScarlet fluorescence was measured every 10 minutes. Curves show the average of three technical replicates; error bars indicate standard deviation. The relative fluorescence (the basis to calculate FS probabilities) was calculated as the ratio of the endpoint fluorescence of the sample and the endpoint fluorescence of 0-control, after subtracting the fluorescence of the negative control (water and reaction mix) (see Methods).

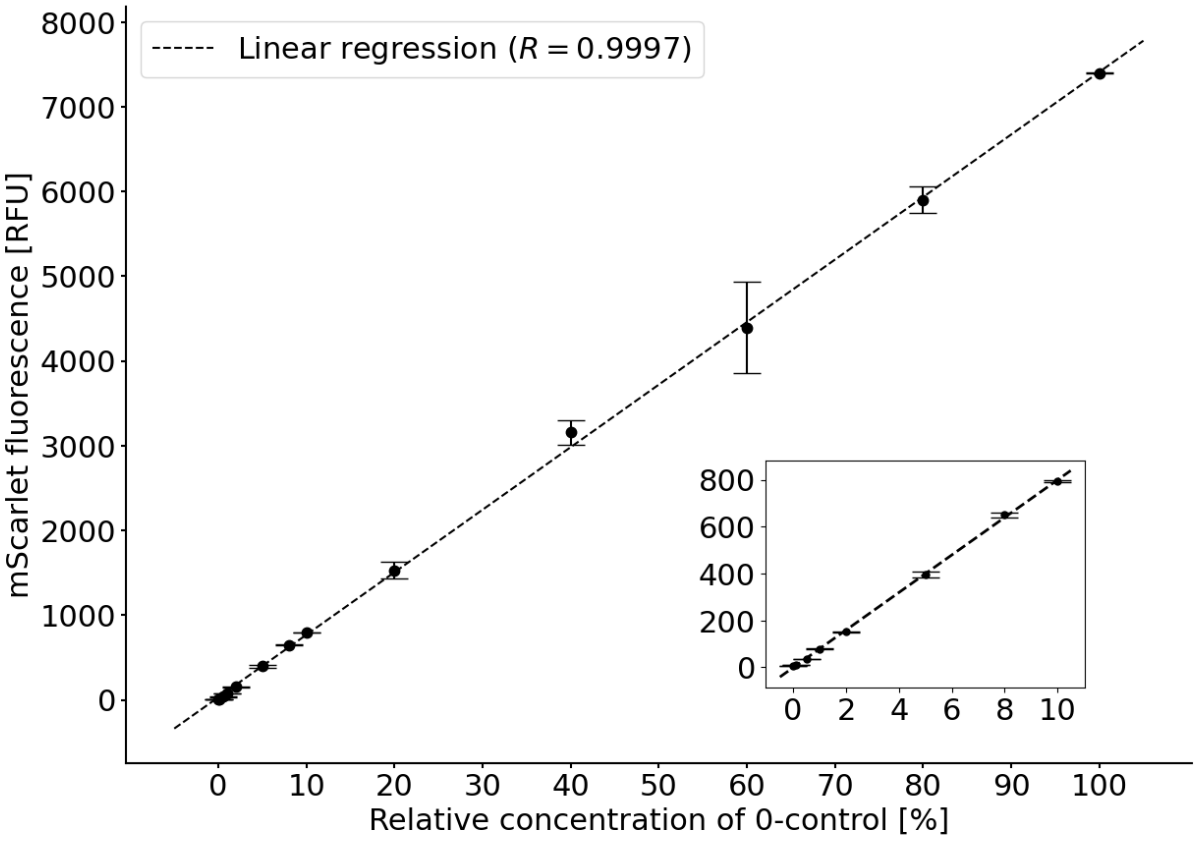


**Figure S3. Linearity of fluorescence detection.**

The transcription/translation reaction of the cell-free expression system with the 0-control plasmid was carried out in the same way as for quantification of frameshifting. The reaction mix was then serially diluted and fluorescence was measured. Relationship between mScarlet-I concentration and fluorescence is linear across the complete range of fluorescence values that were measured for the tested 10-mers: Values for the library ranged from 400 to 2,000 (6,000-8,000 for 0-control).

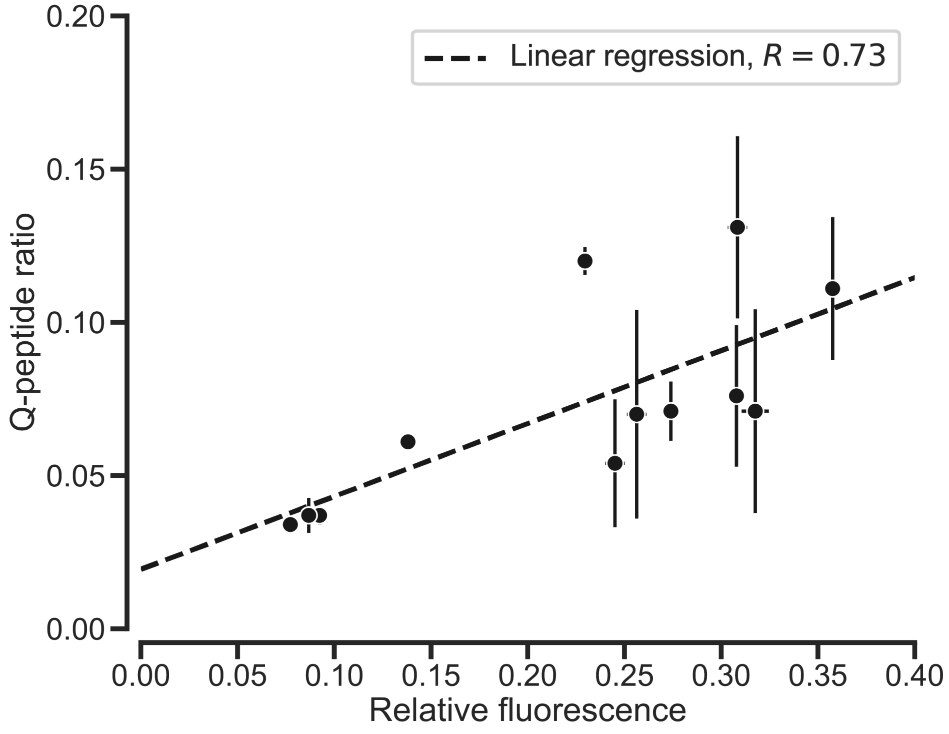


**Figure S4: Quantification of frameshift probabilities by MS (Q-peptide ratio) and fluorescence.**

Frameshift probabilities measured via the median Q-peptide ratio (MS quantification) vs. fluorescence relative to 0-control, measured from the same experiment (error bars represent standard deviation of intensity ratios of the three peptide pairs (y-axis) and standard deviation of three replicate reactions (x-axis)). The correlation between both readouts is good, indicating that the fluorescence signal accurately represents the relative ability of the 10-mers to cause frameshifting, without being strongly affected by different expression levels. Note that raw relative fluorescence was plotted, before applying corrections.

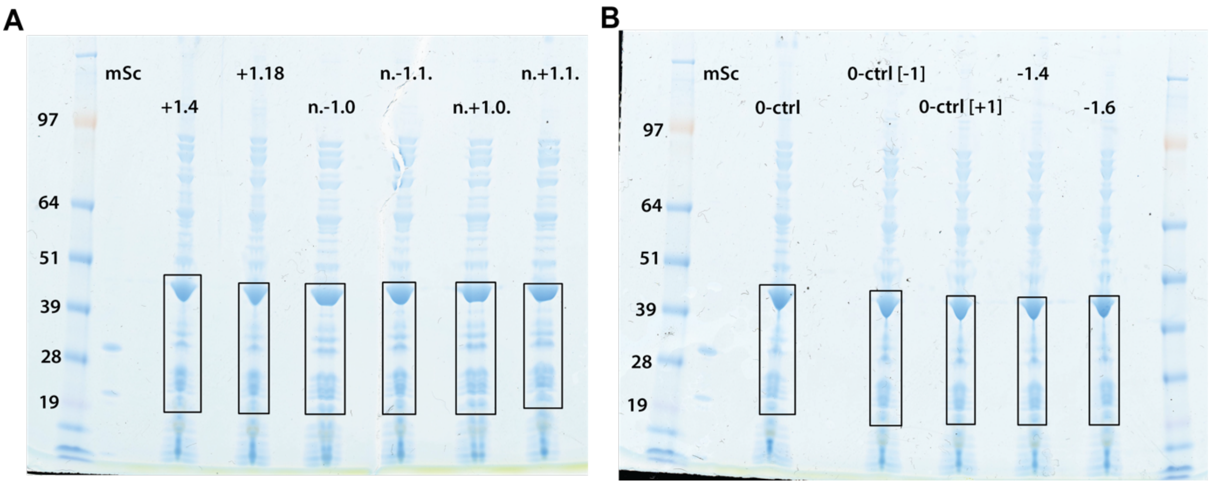


**Figure S5. Selection of molecular weight fraction to quantify alternative initiation.**

Gel used to prepare sample preparation for quantification of alternative initiation. After protein production in the cell-free expression system, samples were loaded onto a gel together with purified mScarlet. The marked gel region corresponds to approx. 19-45kDa and thus contains full-length frameshift product as well as any products of alternative initiation (which would have at least the length of mScarlet), but not products of alternative initiation terminating after the FS site (<10kDa). Q1 peptides were then quantified in the selected fraction.

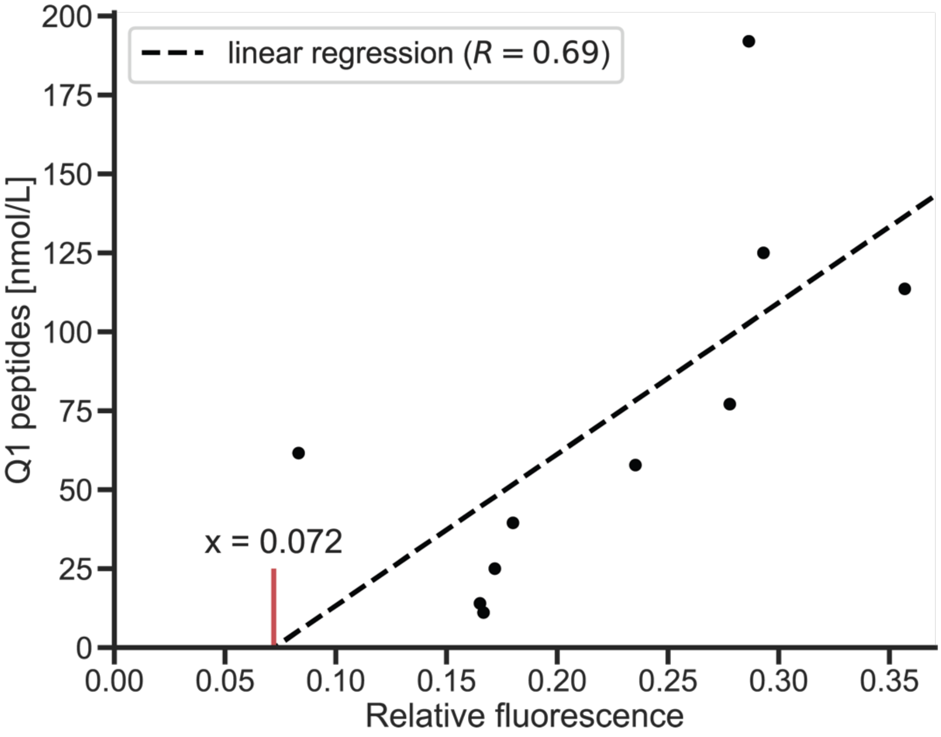


**Figure S6: Determination of fluorescence baseline by absolute quantification of Q-peptides.**

Scatterplot of the molar concentration of Q1 peptides in the protein synthesis reaction mix versus the FS probability as determined by relative fluorescence. The Q1 peptides were quantified from a molecular weight fraction containing only full-length frameshift products and any potential products of alternative initiation (which contain mScarlet, but not Q1, Figure S5). The amount of Q1 and the relative fluorescence correlate well, indicating that alternative initiation causes a relatively fixed baseline of mScarlet/fluorescence, and any extra mScarlet beyond that is part of full-length frameshift products also containing Q1. The intercept of the regression line with the x-axis indicates the amount of mScarlet (relative fluorescence) when no Q1 peptides are present. This amount must then be due to alternative initiation. The intercept with the x-axis (=0.072) is then the baseline relative fluorescence that we used to correct fluorescence-based frameshift probabilities in subsequent analyses.

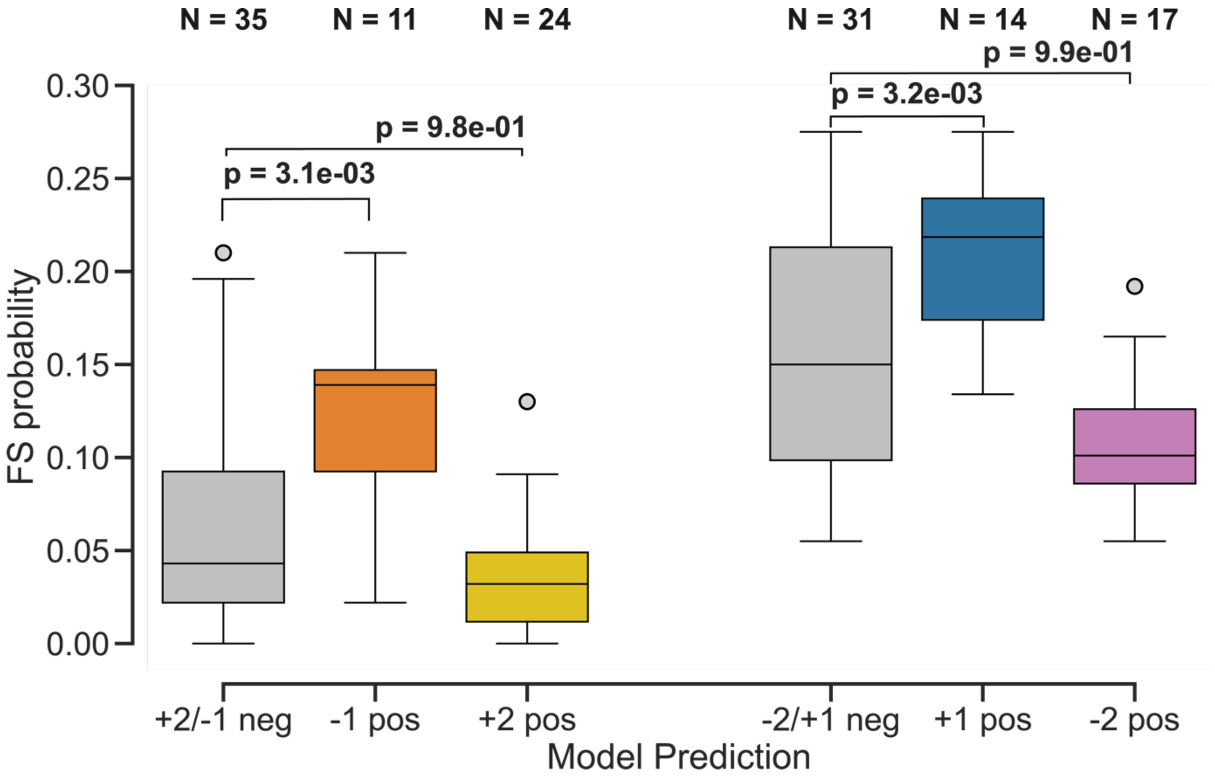


**Figure S7. Measured frameshift probabilities stratified by SLIPPERRS predictions.** 10-mers are classified by frame (left: -1, right: +1) and by the SLIPPERRS prediction. P-values are based on one-sided independent t-test. Since the assay only measures the amount of shifting into a given frame (-1/+1) but not the shift type, -1/+2 and -2/+1 shifts are grouped together. Gray boxes contain sequences for which SLIPPERRS does not predict either of the two shift types for the given frame. While sequences predicted to shift by -1 or +1 show significantly higher FS probabilities than negative predictions, while this does not hold for predicted -2/+2 positives.

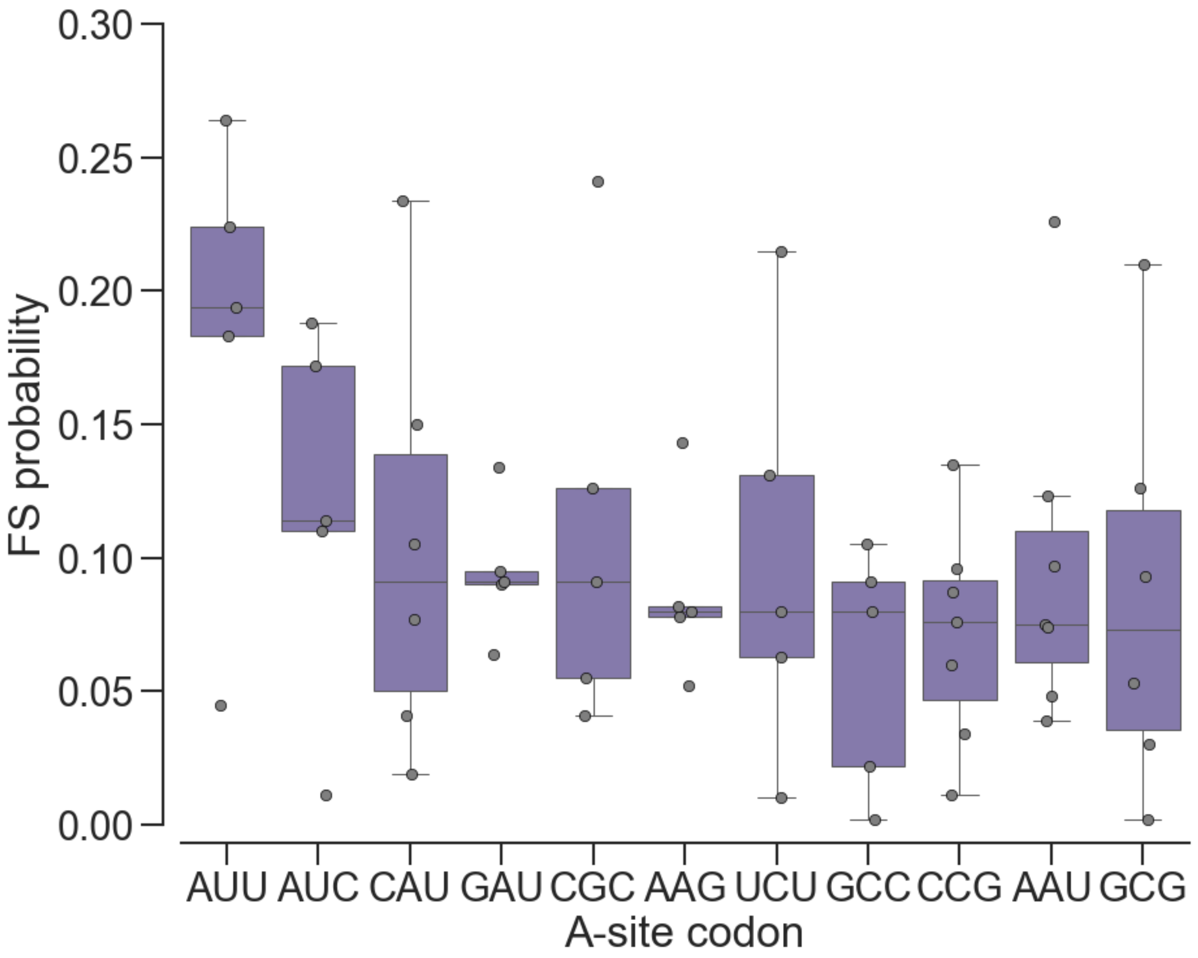


**Figure S8. Frameshift probability by A-site codon.**
Experimentally tested 10-mers are grouped by their A-site codon. Only groups with at least five sequences are shown. The codon groups are sorted by their median frameshift probability.

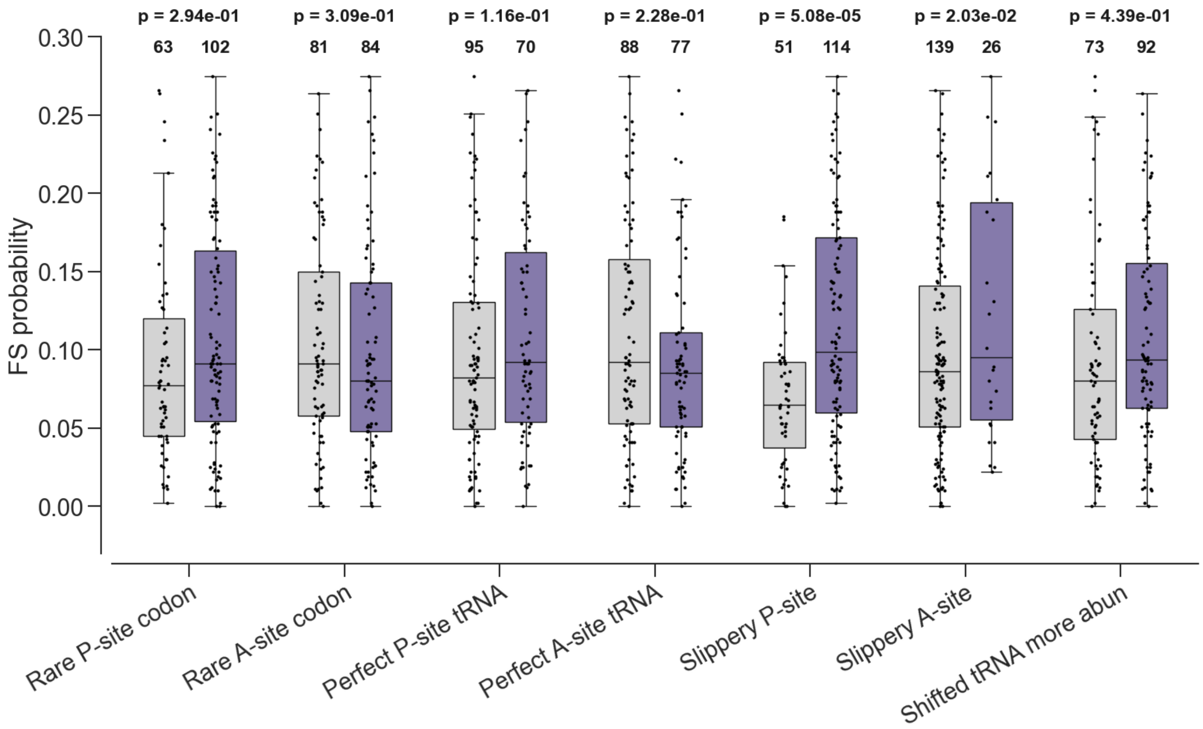


**Figure S9. Influence of 10-mer properties on frameshift probability.**
10-mers were tested for properties associated with frameshifting, such as usage of rare codons or the ability of the 10-mer to allow tRNA rebinding (see Methods for details). Boxplots show the distribution of FS probabilities without (gray) or with (purple) the respective property. P-values were calculated using two-sided independent t-test. Among the investigated sequence properties, a slippery P-site (and codon patterns) is strongly associated with higher frameshift probabilities, and a slippery A-site to a lesser degree.

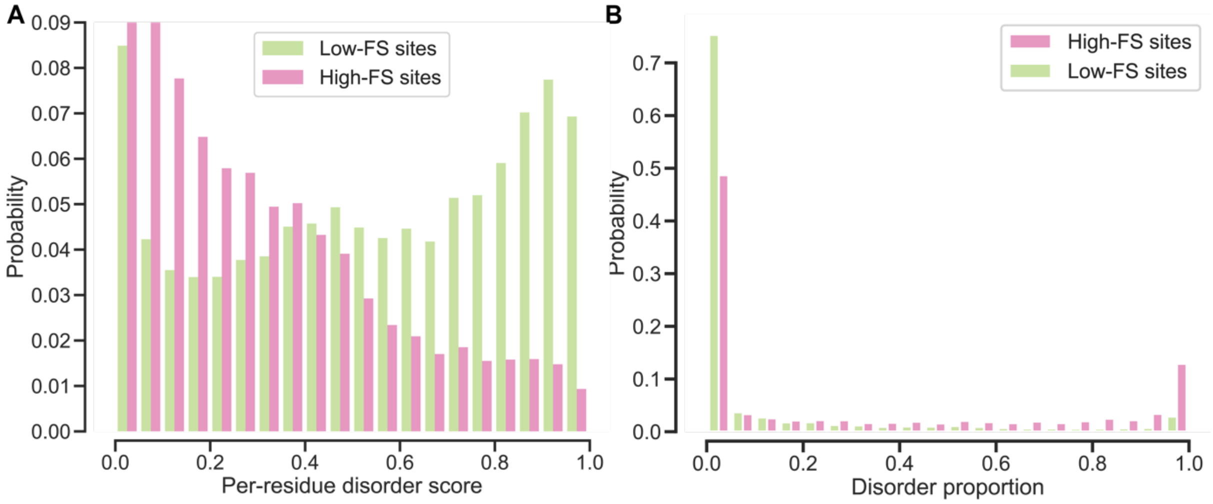


**Figure S10: Disorder content of hypothetical frameshift proteins.**

A. Histogram of IUPred disorder scores for hypothetical frameshift proteins in *Gammaproteobacteria* for low-FS (N =20,253) and high-FS (N = 11,212) sites. Only the sequence region after the FS site was considered, and only sequences where this region was at least 10 residues long, excluding fusions. Bin size is 0.05. Frameshift products of low-FS sites have much higher disorder content than products of high-FS sites.

B. Histogram of per-protein disorder proportion (proportion of sequence with IUPred score >= 0.5) of post-FS regions of hypothetical frameshift proteins for low-FS (N =20,253) and high-FS (N = 11,212) sites. Only the sequence region after the FS site was considered, and only sequences where this region was at least 10 residues long, excluding fusions. Bin size is 0.05. Frameshift products of low-FS sites have much higher average disorder scores than products of high-FS sites.

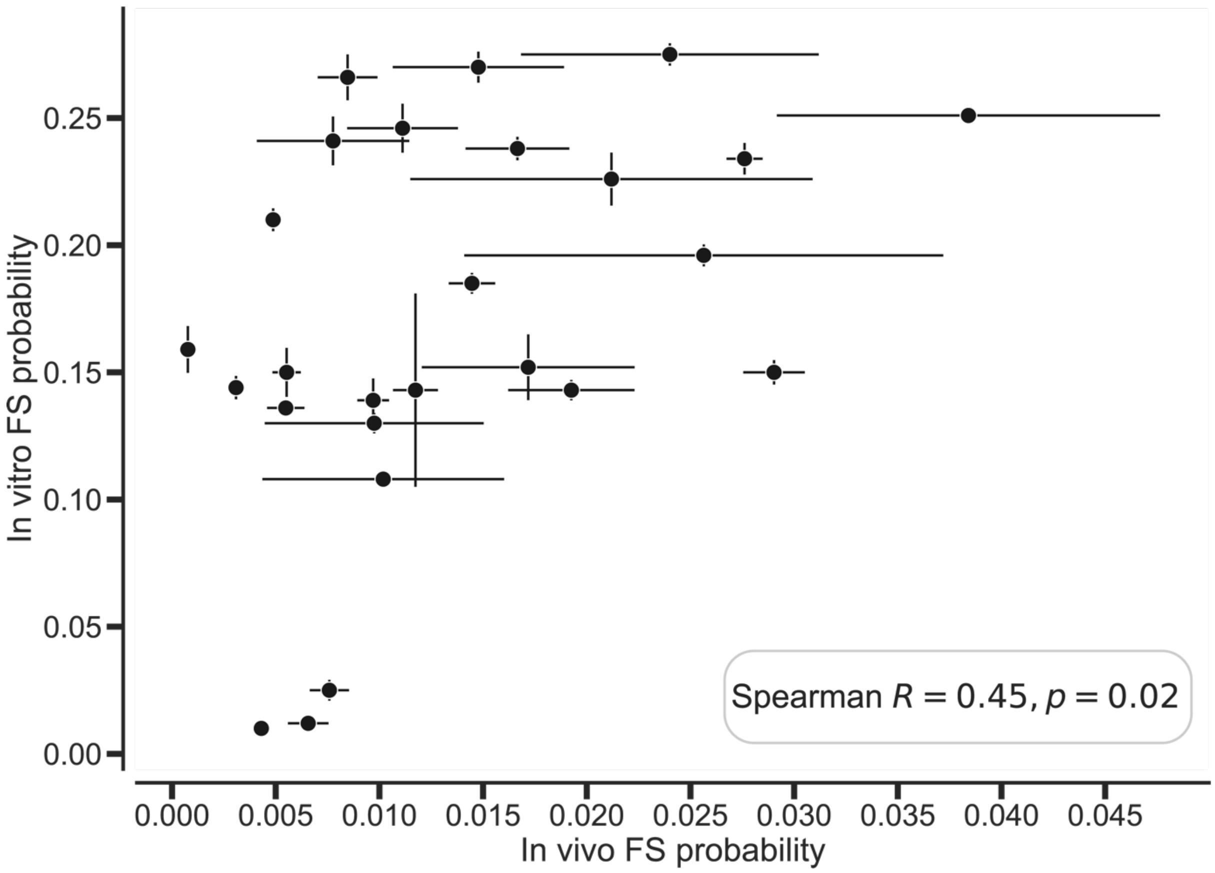


**Figure S11: In cell frameshift quantification correlates with in vitro measurements but it is less sensitive.**

*E. coli* cells were transformed with plasmids containing the same frameshift testing constructs used in the cell-free assays. After 16h of growth, mScarlet fluorescence was measured from living cells. Analogous to *in vitro* measurements, FS probability was calculated as the fluorescence relative to 0-control. *In vivo* and *in vitro* measurements correlate although probabilities measured *in vivo* are much lower and low fluorescence values tend to be indistinguishable, likely due to reduced sensitivity for fluorescence in living cells.

### SUPPLEMENTARY TABLES

**Table S1. Nucleotide interaction parameters for *E. coli.*** One parameter for each combination of codon nucleotide and anticodon nucleotide. L is the modified nucleotide lysidine common in E. coli anticodons. Lower parameters indicate better binding. Parameters in code-compatible input format are provided as supplementary data. Parameters are adapted from Landerer et al. (2024)[1].

| **Codon nucleotide** | **Anticodon nucleotide** | **Parameter** |
|---|---|---|
| A | A | 3.214726 |
| A | C | 3.748255 |
| A | G | 4.411252 |
| C | A | 4.563604 |
| C | C | 8.467473 |
| C | U | 6.019738 |
| T | C | 5.372364 |
| T | U | 8.116570 |
| T | G | 1.054184 |
| G | A | 3.283392 |
| G | U | 0.907996 |
| G | G | 5.132492 |
| A | U | -3.200599 |
| T | A | -2.638884 |
| G | C | -4.155617 |
| C | G | -0.612063 |
| A | L | -8.186579 |
| C | L | -4.894885 |
| T | L | -4.657662 |
| G | L | -1.421939 |

**Table S2. Position parameters for E. coli.**

One parameter for each of the three codon/anticodon positions. Parameters were fitted with the parameter for the third position fixed at 1, such that the parameters for positions 1 and 2 are relative to 1. Parameters in code-compatible input format are provided as supplementary data. Parameters are adapted from Landerer et al. (2024)[1].

| Codon position | Parameter |
| --- | --- |
| 1 | 2.213301 |
| 2 | 1.558439 |
| 3 | 1 |

**Table S3. Quantitative peptides (Q peptides) in the frameshift construct.** For each peptide, all charge states that were used for quantification are given with their respective mass-to-charge ratio (m/z).

| **Name** | **Sequence** | **M/z** | **Charge** |
|---|---|---|---|
| **BSA.1** | HLVDEPQNLIK | 653.3617 | 2 |
| **BSA.1** | HLVDEPQNLIK | 435.9102 | 3 |
| **BSA.2** | LGEYGFQNALIVR | 740.4014 | 2 |
| **BSA.3** | YLYEIAR | 464.2504 | 2 |
| **BSA.4** | DAFLGSFLYEYSR | 784.3791 | 2 |
| **Q1.1** | HLVEEPNQLIK | 660.370361 | 2 |
| **Q1.1** | HLVEEPNQLIK | 440.582855 | 3 |
| **Q1.2** | LGDYGFNNALIVR | 726.386536 | 2 |
| **Q1.3** | YLYDVAR | 450.235535 | 2 |
| **Q1.4** | DAFIGTFLYEYSR | 791.383667 | 2 |
| **Q2.1** | HVLEEPQQLIK | 667.378174 | 2 |
| **Q2.1** | HVLEEPQQLIK | 445.254730 | 3 |
| **Q2.2** | LGDYGFNNLVIVR | 740.402161 | 2 |
| **Q2.3** | YLDYAAR | 436.219879 | 2 |
| **Q2.4** | DAFIGTFYLDYSR | 784.375793 | 2 |
| **mScarlet.1** | LDITSHNEDYTVVEQYER | 1106.014526 | 2 |
| **mScarlet.1** | LDITSHNEDYTVVEQYER | 737.678894 | 3 |
| **mScarlet.2** | LYPEDGVLK | 517.282654 | 2 |

### References


1. Landerer, C., Poehls, J. & Toth-Petroczy, A. Fitness effects of phenotypic mutations at proteome-scale reveal optimality of translation machinery. *Molecular Biology and Evolution* **41**, msae048 (2024).
